# A hierarchical Bayesian framework accommodates intraspecific and interspecific variation in multivariate traits

**DOI:** 10.64898/2026.05.28.728593

**Authors:** Levi Yoder Raskin, Maja Šešelj, John Huelsenbeck, Wonseop Lim, Jacky Kaiyuan Li, Debbie Guatelli-Steinberg, Mackie C. O’Hara, Bárbara Domingues Bitarello

**Affiliations:** Department of Integrative Biology, University of California, Berkeley; Department of Anthropology, Bryn Mawr College; Department of Biology, Bryn Mawr College; Biostatistics Division, University of California, Berkeley; Department of Anthropology, The Ohio State University; Department of Biology, Ball State University

**Author notes:** **Correspondence:** Levi Yoder Raskin. M.O. and B.B. contributed equally to this work. **Author contributions:** Conceptualization: LYR, MŠ, BDB, MCO, and JH. Formal analysis: LYR, MCO, and DGS. Investigation: LYR,MCO, and DGS. Methodology: LYR, JH,WL, and JL.Resources: JH, BDB, and MŠ. Software: LYR. Supervision: JH, MŠ, and BDB. Validation: LYR. Visualization: LYR. Writing - Original Draft Preparation: LYR. Writing - Review & Editing: LYR, MŠ, JH, WL, JL, DGS, MCO, and BDB.

**Keywords:** phylogenetic comparative methods, intraspecific variation, Bayesian inference, enamel growth, human evolution

## Abstract

Phylogenetic comparative methods are a critical tool in biology, providing the framework to test evolutionary hypotheses of phenotypic diversification. Accommodating intraspecific variation in multivariate analyses is critical for accurate evolutionary inference, but current methods that incorporate intraspecific variation either 1) assume that traits evolve independently or 2) that all taxa share the same intraspecific covariance structure. Violations of these assumptions can produce biased estimates of evolutionary parameters. Here, we introduce a hierarchical Bayesian framework for multivariate traits that jointly estimates taxon-specific intraspecific covariance structures alongside the underlying evolutionary process. This framework propagates uncertainty from sample size discrepancies and missing data, enabling the incorporation of highly variable morphological traits into phylogenetic analyses. Analysis of simulated data confirms that the model and implementation are well calibrated under the assumed generative model, including challenging datasets with more traits than individuals and substantial missing observations. Applied to perikymata spacing across the great ape clade, including modern humans and Neandertals, the framework recovers intraspecific covariance structures that differ among taxa and yields evolutionary rate estimates markedly more uniform across the tooth crown than those obtained when taxon means are fixed. Our method, which is applicable to other multivariate traits, provide a flexible, tractable approach to joint estimation of intraspecific variation and evolutionary process in multivariate traits.

## Introduction

A core goal of evolutionary biology is to characterize the patterns and processes by which organisms diverge. Traits that vary across lineages are especially informative, as they may reflect lineage-specific developmental or evolutionary processes. Phylogenetic comparative methods generally model interspecific variation as the product of evolutionary processes such as Brownian motion or the Ornstein-Uhlenbeck process (Felsenstein, 1985; Hansen, 1997). It is well known that reducing the trait information for multiple individuals of a species to a single estimate (typically an average) can lead to biased estimates of evolutionary parameters (Garamszegi, 2014; Garamszegi and Møller, 2010; Harmon and Losos, 2005; Silvestro et al., 2015). As such, many approaches attempt to accommodate for intraspecific variation in phylogenetic comparative analyses with methods such as mixed models (Adams and Collyer, 2024; Clavel et al., 2015; Felsenstein, 2008; Housworth et al., 2004; Ives et al., 2007; Lynch, 1991; Villemereuil et al., 2012), bias correction (Hansen and Bartoszek, 2012), and Bayesian hierarchical models (Gaboriau et al., 2020; Kostikova et al., 2016; Revell and Reynolds, 2012). For univariate traits, methods have been developed which estimate per-taxon intraspecific distributions and propagate uncertainty in these intraspecific distributions through to the evolutionary parameters (Gaboriau et al., 2020; Kostikova et al., 2016). However, for multivariate traits, existing approaches assume that all species either share the same intraspecific variance-covariance (VCV) matrix (Felsenstein, 2008) or do not explicitly estimate species-specific multivariate intraspecific covariance structures (Adams and Collyer, 2024; Nakagawa et al., 2025).

These limitations are problematic when 1) rapid evolution and reorganization of covariance structure have occurred within a clade, 2) a clade of interest has taxon-specific patterns of morphological covariance, or 3) sample size varies across taxa (e.g., Adams and Felice, 2014). The first two scenarios may result in there truly being multiple intraspecific VCV matrices due to the evolutionary changes that have occurred within a sub-clade or individual taxa, whereas the third scenario results in different levels of statistical uncertainty in the intraspecific VCV across the clade. Even without sample size discrepancies, statistical uncertainty in the intraspecific VCV may vary from taxon to taxon due to specific pattern of missing data within each taxon. Because of these methodological limitations, these traits cannot readily be accommodated by existing multivariate comparative approaches without either imposing a shared within-taxon covariance structure or adopting other restrictive approximations.

Incremental enamel growth traits exemplify the limitations of current multivariate phylogenetic methods and will serve as a case study for a new method we present here. As teeth develop through the incremental deposition of enamel from cusp to cervix, the process leaves behind long-period striae (Risnes, 1998). When these striae emerge on the surface of the tooth, they are known as perikymata. Within a tooth crown, regions with more perikymata per unit area indicate a slower local enamel growth rate than those with fewer perikymata per unit area. Characterizing the distribution of perikymata across a crown is known as the perikymata packing pattern and, while each tooth varies, species have noticeably different trends in their perikymata packing patterns (Dean and Reid, 2001; Guatelli-Steinberg et al., 2018). For example, humans (*Homo sapiens*) are known to have extreme perikymata distributions, with tightly packed perikymata in the cervical half of the tooth (Figure 1; Dean and Reid, 2001; Guatelli-Steinberg et al., 2018; Xing et al., 2015). While perikymata have long been interpreted as informative for hominin taxonomic comparisons (e.g., García-Campos et al., 2022; Guatelli-Steinberg et al., 2018; Modesto-Mata et al., 2020, 2022; Xing et al., 2015), there has never been a phylogenetic analysis of this incremental dental growth feature in the great apes. Due to the inherently multivariate and highly correlated nature of perikymata, the degree of interspecific variation with clear taxon-specific patterns within the great ape clade, and the likewise large amount of intraspecific variation within great ape taxa (Dean and Reid, 2001; Guatelli-Steinberg et al., 2018), perikymata are a prime case study for a multivariate phylogenetic comparative analysis that estimates taxon-specific patterns of intraspecific variation.

**Figure 1:**
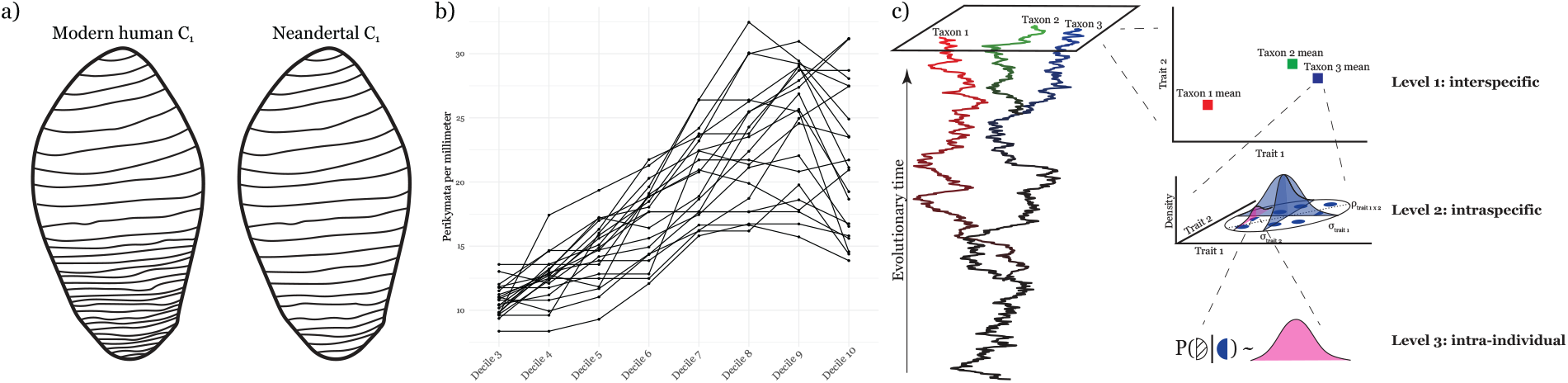
Perikymata schematic and depiction of modern human variation. a), a schematic representation of perikymata distribution across modern human and Neandertal mandibular canine (C_1_) is depicted. Note that modern humans typically exhibit denser perikymata towards the cervix (bottom) than the cusp (top), while Neandertals exhibit a less drastic increase. b), modern human C_1_ perikymata per millimeter per decile. Each line represents the perikymata per millimeter per decile for an individual modern human. c), a graphical depiction of our modeling scheme. Level 1: interspecific variation, modeled with a multivariate Brownian motion. Level 2: intraspecific variation, modeled with a multivariate normal distribution. Level 3: intra-individual variation, missing data modeled with a conditional normal distribution, incorporating both what observations are preserved for that individual and population-level data in estimation of the missing element.

Here we present a hierarchical Bayesian framework that estimates taxon-specific intraspecific covariance structure jointly with the evolutionary process; validate it by simulation across a range of trait dimensionalities, incompleteness, and tree sizes; and apply it to perikymata spacing across the great ape clade. This framework allows for multivariate traits with taxon-specific patterns of intraspecific variation, such as perikymata, to be analyzed with taxon-specific intraspecific covariance structure for the first time. Through a Bayesian approach, we propagate uncertainty across each level of the hierarchy, from interspecific to intraspecific to intra-individual. This allows us to assess whether apparent differences in perikymata spacing are truly a result of evolutionary processes or an artifact of intraspecific variation and, potentially, sampling (Figure 1).

The approach developed here can be used for any multivariate trait (e.g., any complex phenotypic trait, be it morphological, behavioral, physiological, or related to life history), both within and beyond the great ape clade. In particular, our methods can shed light on evolutionary patterns of traits that are highly variable within taxa, but require a sufficiently rigorous statistical approach that can appropriately accommodate unique intraspecific variance-covariance structures for phylogenetic comparative analyses across taxa. We provide an accessible command line software implementation of this new method written in C++, called BURL: Between and within-group Uncertainty in Rates across Lineages, available at: https://github.com/Levi-Raskin/PerikymataPhylogenetics/.

## Materials and Methods

### The model

#### Definitions and Notations

Let *X*_*i*_ refer to the ℝ^*n×d*^ data at taxon *i*, where *d* is the number of traits sampled from *n* individuals. For example, in our application towards perikymata, *X*_Neandertal_ refers to the perikymata per millimeter per decile (PMD) data for each of seven Neandertals from whom we have C_1_ (lower canine) data. In this case, *n* = 7 and *d* = 8 (the number of deciles considered). We assume that the true mean for taxon *i*, denoted *µ*_*i*_ *∈* ℝ^*d*^, (*d* free parameters), evolves according to a multivariate Brownian motion process on a phylogeny Ψ with an evolutionary VCV, denoted as 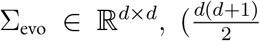 free parameters). In our analyses, Ψ is taken to be fixed and known without error, but this assumption can be relaxed. Let *m* denote the number of tips in the phylogeny, taken here to represent species. We assume that each individual at each tip is drawn from a normal distribution with a tip-specific VCV matrix, denoted as 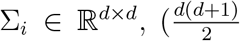 free parameters), and the tip-specific mean (*µ*_*i*_). For convenience, let *S* indicate all the Σ_*i*_, *M* indicate all the *µ*_*i*_, and *X* indicate all the *X*_*i*_. Note that all *S* and Σ_evo_ are constrained symmetric positive definite.

#### Jointly modeling inter- and intraspecific variation

We model intraspecific variation for the *i*th tip as

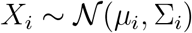

and interspecific variation (the evolutionary component) as

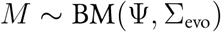

The joint posterior probability distribution of the parameters for our model is:

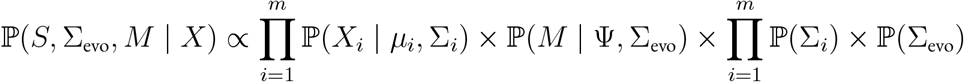

We note that the multivariate Brownian Motion model acts as an informed prior distribution on *M*.

#### Inference under the model

We are able to draw directly from the posterior distribution for each Σ_*i*_ given *X*_*i*_, *µ*_*i*_ as follows:

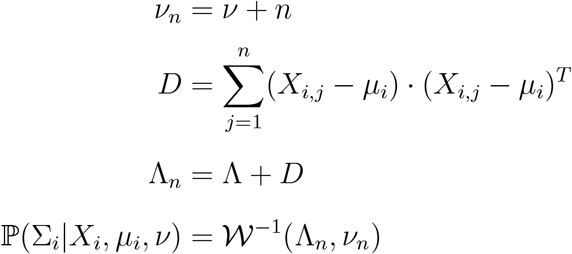

Here, *X*_*i,j*_ denotes the ℝ^*d*^ vector containing all the observations from individual *j* in species *i*. We there-fore draw our new estimate for Σ_*i*_ directly from ℙ (Σ_*i*_|*X*_*i*_, *µ*_*i*_, *ν*).

Inspired by Lartillot and Poujol (2011), we implement a Gibbs sampler for Σ_evo_. First, we calculate Felsenstein’s standardized independent contrast following Felsenstein (1985). These standardized independent contrasts are stored in a ℝ^(*m*−1)*×d*^ matrix, denoted *Y*. Note that *Y*_*j*_ ~ *N* (**0**, Σ_evo_). We can then draw Σ_evo_ from the posterior distribution as follows:

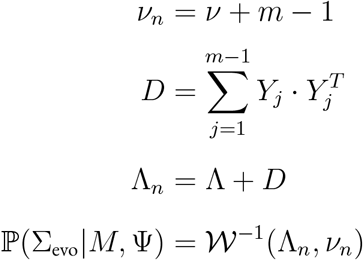

By choosing an inverse Wishart prior for *S* and Σ_evo_, we guarantee by construction that every Σ_*i*_ in our posterior distribution will be non-singular and appropriately symmetric positive definite even if a given tip has many fewer individuals than traits. This construction circumvents known issues of maximum likelihood inference of variance-covariance matrices when *d* ≥ *n* (Clavel et al., 2019). *We place the inverse Wishart prior on Σ*_*evo*_ *and on each Σ*_*i*_, with *ν* = *d* + 2 and a scale matrix Λ having 1 on the diagonal and 1 *×* 10^−6^ off the diagonal. We chose *ν* = *d* + 2 because it is the smallest integer degrees of freedom for which the inverse Wishart mean is defined; the prior is therefore centered on E[Σ_*i*_] = E[Σ_evo_] = Λ, that is, on unit variances and negligible covariances, and is the least informative proper choice within this family.

We capitalize on the above modeling structure to impute missing data, common across morphological datasets, but especially for fossil taxa. By using conditional normal distributions, for each individual with missing data, we can impute the missing traits by drawing directly from the posterior distribution: a normal distribution conditioning on the observed traits for that individual, *µ*_*i*_, and Σ_*i*_. We note that *X*_*i*_ used in the Gibbs update for Σ_*i*_ is this completed data matrix, combining observed and imputed values.

Unfortunately, a Gibbs update for *µ*_*i*_ is nontrivial as we need to condition not just on *X*_*i*_ and Σ_*i*_, but also on Ψ, Σ_evo_, and *µ*_*j i*_. Instead, we rely on the Metropolis-Hastings algorithm to sample from this posterior distribution (Hastings, 1970; Metropolis et al., 1953). Proposals are drawn from a multivariate normal distribution centered on the current *µ*_*i*_ with 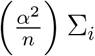 where *α* is adaptively tuned for the first 50,000 times each *µ*_*i*_ is proposed. All samples drawn from the posterior distributions are taken after this adaptive tuning has completed.

### Simulation study design

We conducted a simulation study to assess whether our model and inferential scheme were well-calibrated across a range of tree sizes, trait sizes, completeness, and ratios between *d* and *n*. We simulated datasets with conditions varying in four ways: number of taxa (8, 16, and 32); number of traits (8, 16, and 32); number of individuals per taxon (2, 4, 8, and 16); and number of missing observations (i.e., individual *i* is missing trait *j*) across the entire dataset (0, 8, 16, and 32). For each unique condition (e.g., 8 taxa, 16 traits, 2 individuals per taxon, no missing data) we sampled 100 sets of parameters from the prior and simulated a dataset under those parameters. We then ran 10,000,000 cycles of Markov chain Monte Carlo (MCMC), a standard numerical method to estimate complex probability distributions, sampling every 1,000 cycles and calculated how often the sampled parameters were in our 95% credible interval. For matrix-valued parameters (*S*, Σ_evo_), we assessed element-wise marginal intervals. This procedure enabled us to assess the accuracy of our model for datasets across a range of challenging conditions, including poorly identified *S* where *d* : *n* = 32 : 2, poorly identified Σ_evo_ where *d* : *m* − 1 = 32 : 7, and a dataset incompleteness percentage of up to 25%. We note that since truths are drawn from the prior, this simulation study only assesses computation under a correctly specified model, not robustness to prior or model misspecification in empirical applications. Further, under the same simulation scheme, we compare the coverage of the full model to one where we fix species means: 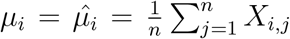. This comparison assesses the importance of accounting for intraspecific variation. For the taxon means model only, we ensured that all simulated datasets contained at least one individual observed per trait (i.e., so that if there are two individuals in a taxon, both cannot be missing trait *i*); for the full model, no such restriction was applied.

In addition to this coverage test, which evaluates a binary inclusion/exclusion across 144 conditions, we used simulation-based calibration (Cook et al., 2006; Modrák et al., 2025; Talts et al., 2020) to confirm that our samplers for *S*, Σ_evo_, and *M* inferred correctly-shaped posterior distributions. We assessed four different simulation conditions, all with eight taxa and eight traits: 0 or 32 missing observations across the entire dataset and 2 or 16 individuals per taxon. For each condition, we sampled 5,000 sets of parameters from the prior and simulated a dataset under those parameters. We then ran 10,000,000 cycles of MCMC and calculated the normalized rank in the inferred posterior distribution of the trace of true values of *S* and Σ_evo_ and the element-wise true values of *M* (see Cook et al., 2006; Talts et al., 2020, for more detail). These normalized ranks should be uniformly distributed; to test for uniformity, we apply the *γ* statistic of Säilynoja et al. (2022), where log(*γ/γ*_5_) *<* 0 indicates rejection of uniformity at the 5% significance level. We used the R package *bayesplot* to calculate *γ*_5_ (Gabry et al., 2019). Reducing *S* and Σ_evo_ to traces allows for coherent ranking of these parameters, but this dimensionality reduction obscures miscalibration concentrated off the diagonals. However, as *S* and Σ_evo_ are updated exactly through Gibbs draws from their conditional posterior distribution, calibration failures in *S* and Σ_evo_ can only enter through *M*; thus, the trace of these matrices is sufficient to confirm that the sampler for *M* is not biasing, narrowing, or otherwise impacting sampling from *S* and Σ_evo_.

All analyses in the simulation study were performed on a server containing two 64-thread Intel Xeon 6737P processors with a base clock frequency of 2.9 gigahertz. Mean wall clock time for each of the 144 simulation conditions tested in the full coverage test is reported.

### Empirical application

#### Dataset

We used perikymata count per decile and reconstructed cusp height (RCH) data from modern humans (C_1_: *n* = 21, I^2^: *n* = 40), Neandertals (C_1_: *n* = 7, I^2^: *n* = 9), *Pan paniscus* (C_1_: *n* = 32, I^2^: *n* = 2), *Pan troglodytes* (C_1_: *n* = 14, I^2^: *n* = 6), *Gorilla beringei* (C_1_: *n* = 11), *Gorilla gorilla* (C_1_: *n* = 8), *Pongo abelii* (C_1_: *n* = 4), and *Pongo pygmaeus* (C_1_: *n* = 13). These totals include previously-published modern human I^2^ perikymata spacing data from Modesto-Mata et al. (2017) (*n* = 4) and previously-published Neandertal I^2^ perikymata spacing data from Hlusko et al. (2013) (*n* = 4). There are 165 of 880 C_1_ and 5 of 456 I^2^ observations missing. These great ape and hominin taxa were selected because of accessible perikymata count and RCH data, as well as genomic data. Thus, we can capitalize on a well-resolved, time-calibrated phylogeny (Arnold et al., 2010) on which we can model perikymata count data. While we take the phylogeny used here to be fixed, in other clades where the phylogeny is less certain this assumption could be relaxed. Because perikymata count per decile data is ordinal, we scaled perikymata per decile by the height of each decile 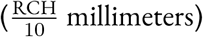, which allows informed application of standard continuous phylogenetic models like multivariate Brownian motion to these data. We excluded decile one and two from analysis, as perikymata counts for these deciles are typically affected by wear and are thus hard to estimate with sufficient accuracy from surface dental casts. Worn crowns were digitally reconstructed to their estimated unworn height using methods established by Saunders et al. (2007) prior to counting perikymata per millimeter per decile of the estimated unworn height. This scaling also accounts for size variation among taxa. For both the C_1_ and I^2^ analysis, we ran 100 million MCMC cycles, sampled every 1,000 cycles, and discarded the first 10% as burn-in. In total, for the C_1_ analysis including hominins, we inferred 553 free parameters, and for the I^2^ analysis we inferred 217.

#### Convergence and model fit

We confirmed convergence by calculating the rank-normalized Gelman-Rubin statistic across four independent analyses as implemented in the R package *posterior* (Vehtari et al., 2021). Conditional on *M*, each Σ_*i*_ and Σ_evo_ posterior is exactly inverse Wishart; marginally, each is a mixture of inverse Wisharts over *M* and the imputed observations. We therefore fit a single inverse Wishart to each marginal posterior as a tractable approximation for computing KL divergence from the prior and symmetrized KL divergences among posterior distributions (*D*_*KL*_(*A*∥*B*) + *D*_*KL*_(*B*∥*A*), Kullback and Leibler 1951). We report the fitted degrees of freedom alongside each divergence as an indication of how heavy-tailed each mixture is. All code for this maximum likelihood estimation and KL divergence calculation was written in R.

Additionally, we leverage the posterior predictive distribution to assess the fit of our model to both the C_1_ and I^2^ datasets with a posterior predictive check. The posterior predictive distribution is the probability describing unobserved individuals for each species; by drawing from the posterior predictive distribution we are, in essence, describing the PMD distributions for the taxon writ large. A posterior predictive check makes repeated draws from the posterior predictive distribution and compares a summary statistic, typically mean and standard deviation, against the original dataset (Gelman et al., 1996; Rubin, 1984). For each posterior sample, predictive draws are generated from a multivariate normal distribution. We randomly sub-sampled without replacement 10,000 posterior samples from our full posterior distribution to reduce the computational cost while retaining a representative sample of the posterior. For each sampled set of parameters, we made 100 draws from the posterior predictive distribution per taxon. This procedure gives a posterior predictive distribution from which taxon-level posterior predictive means and standard deviations can be calculated. These posterior predictive means and standard deviations are then compared against the observed taxon means and standard deviations per decile. We used the Bayesian *p*-value, the percentage of posterior predictive summary statistics greater than or equal to the empirical summary statistic, to assess the fit of our model to our dataset (Gelman et al., 1996; Rubin, 1984). Bayesian *p*-values close to 0 or 1 indicate the model poorly captures that aspect (either mean or standard deviation) of the original dataset.

To test whether our perikymata dataset truly supports unique intraspecific VCV matrices for each taxon, we calculated the symmetrized KL divergence between the intraspecific VCV posterior distributions inferred for each pair of taxa within a genus. Symmetrized KL divergence is zero only when two posterior distributions are identical and grows with their dissimilarity; because it has no absolute scale, we interpret these divergences comparatively across pairs rather than against a fixed threshold. To our knowledge, no available implementation jointly estimates a full, taxon-specific intraspecific covariance matrix for every tip alongside the evolutionary rate matrix while propagating uncertainty from missing data, and none does so in a regime where *d* ≥ *n* for several tips. The closest available comparator instead assumes this structure is shared across taxa, equivalent to the Felsenstein (2008) model as implemented in the R package *rphylopars* (Goolsby et al., 2017). *Thus, we used rphylopars* to calculate the maximum likelihood evolutionary and singular intraspecific VCV matrices under the Felsenstein (2008) model and the R package *RiemBase* (You, 2025) to calculate geodesic distances on the symmetric positive definite manifold (a measure of dissimilarity between VCV matrices that respects their mathematical structure) between these estimates and our posterior distributions for Σ_evo_ and each of the Σ_*i*_. These analyses characterize 1) the extent to which posterior distributions for taxon-specific covariance matrices differ, 2) whether these covariance estimates occupy similar regions of SPD space, and 3) how the inferred evolutionary covariance differs from that obtained under a shared intraspecific covariance assumption.

We assessed statistical uncertainty in the VCV posterior distributions through application of Fréchet variance. Fréchet variance generalizes the notion of variance to non-Euclidean spaces:

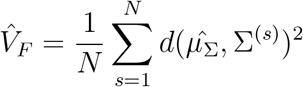

where *N* is the number of samples in our posterior distribution,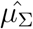 is the Fréchet mean of those posterior samples on the SPD manifold, and 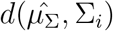 is the geodesic distance relating each sample to that mean (Dubey and Müller, 2019; Fréchet, 1948). We used the *rbase*.*mean* function from the *RiemBase* R package (You, 2025) to calculate the empirical mean VCV matrix on the SPD manifold for the evolutionary and intraspecific VCV parameters in both tooth types. We used this empirical mean VCV to estimate the Fréchet variance of our posterior distribution, using the affine-invariant geodesic distance on the SPD manifold as our metric. Larger Fréchet variance indicates greater uncertainty in the posterior distribution. Quantifying the uncertainty in the mean posterior distributions is simpler than for the VCV parameters as the means are Euclidean. As such, ordinary variance is well-defined for these parameters, and we summarize posterior uncertainty in the means as the total posterior variance across all deciles.

#### Biological interpretation of posterior distributions

Because our primary empirical interest lies in the full distribution of PMD within each taxon while accounting for both intraspecific variance and uncertainty in our parameter estimates, we utilized the posterior predictive distribution to summarize these joint effects. We made a single draw for each taxon from each sample we took from the posterior distribution using the R package *MASS* (Venables and Ripley, 2002) in R v4.4.1. Additionally, we used the R package *overlapping* for calculating overlap between intraspecific distributions (Pastore, 2018). Additionally, we used *ggplot* (Wickham, 2016), *gghalves* (Tiedemann, 2022), *and ggtree* (Yu et al., 2017) *for visualization*.

## Results

We inferred 1) the evolutionary VCV matrix, 2) a unique intraspecific VCV matrix for each taxon, 3) the species mean perikymata per millimeter per decile (PMD), and 4) the missing PMD. These four types of parameters were combined across three hierarchies: 1) the evolutionary level, where we modeled the evolution of the species mean PMD through a multivariate Brownian motion with the evolutionary VCV; 2) the intraspecific level, where we modeled variation among individuals at each species as a multivariate normal distribution with the species mean and intraspecific VCV as parameters; and 3) the intra-individual level, where we modeled the missing data for a given individual through a conditional multivariate normal distribution with the species mean, intraspecific VCV, and observed values for that individual as parameters (Figure 1). This layering mimics the truism that individuals do not evolve, populations do.

### Inference is well-calibrated across diverse and challengingly parameterized datasets

To validate this model and our implementation, we tested the coverage probability for parameters inferred from simulated datasets under the full model that accounts for intraspecific variation. We expect the true value under which we simulated to be in the 95% credible interval in 95% of simulations and, for our full model, our coverage matches expectations across all combinations of parameters with deviations well within expected Monte Carlo error (Figure 2a, individually listed in SOM Table S1). In contrast, we found that, when intraspecific variation exists in a dataset, aggregation of individuals into a single mean dramatically reduces the coverage (Figure 2a). This simulation study also presented an opportunity to characterize model scaling with number of individuals per taxon, number of traits, and number of taxa. While this model, and our implementation of it, is exceptionally fast—10 million cycles of MCMC on average across all parameters take less than 20 minutes running serially—scaling is worst with increasing number of traits, while increasing the number of individuals per taxon and increasing the number of taxa have substantially weaker effects on run time (Figure 2b). Additionally, our rank uniform histograms are all appropriately uniform with no more deviations from uniformity than would be expected given *α* (Figure 2c, SOM Figures S27-29). Taken together, our model and implementation is well-calibrated and performant with tractable run times.

**Figure 2:**
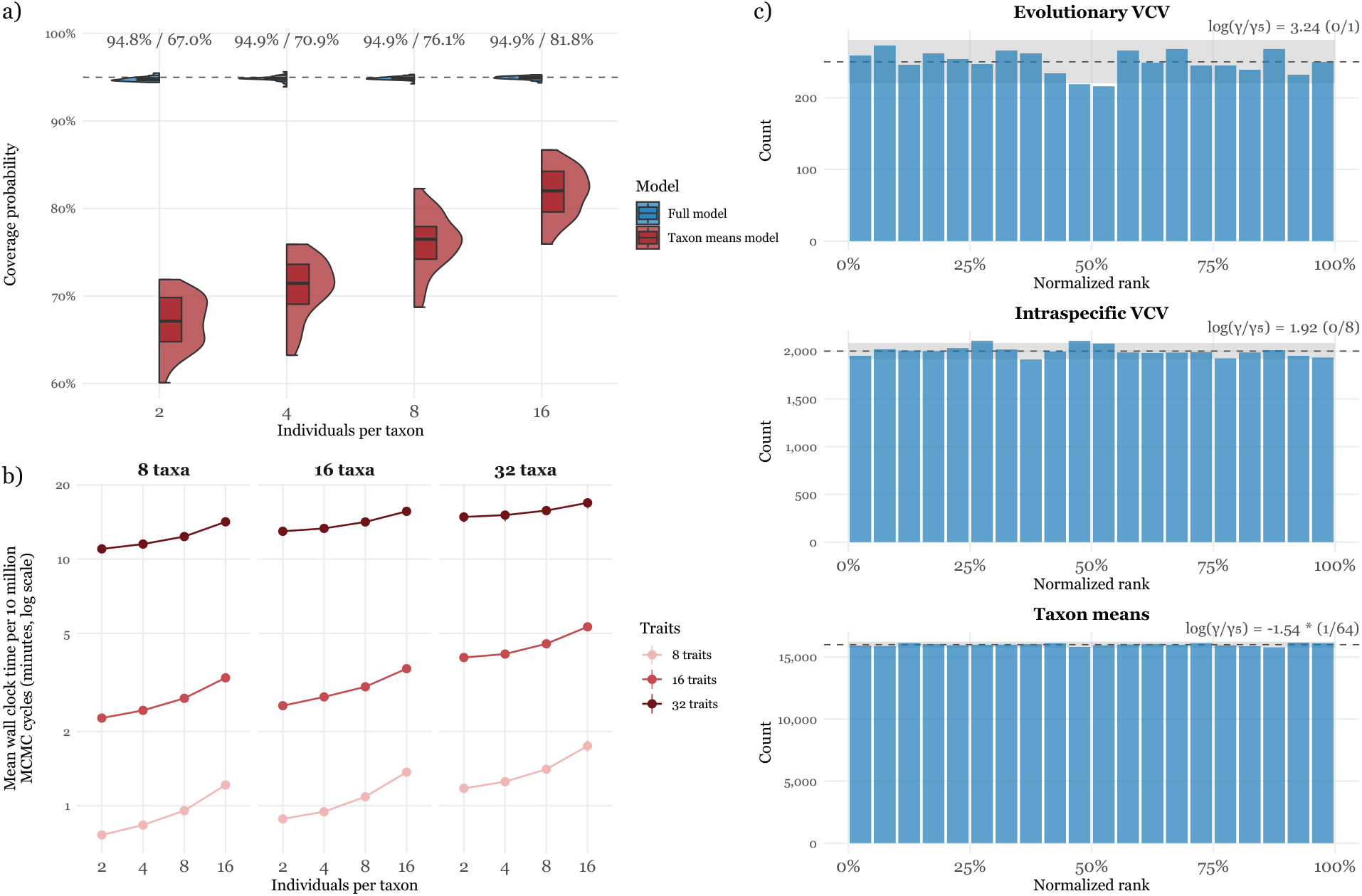
Simulation-based validation and computational performance of BURL. a) Empirical coverage of nominal 95% marginal credible intervals for the full hierarchical model and the fixed-taxon-means procedure. Points show mean coverage at each number of individuals per taxon, averaged across the remaining simulation factors; condition-specific results are reported in SOM Table S1. b) Mean wall-clock time per 10 million MCMC cycles as a function of the number of individuals per taxon and number of traits. Values are averaged across the number of missing observations. c) Simulation-based calibration results for a representative condition with eight taxa, eight traits, 16 individuals per taxon, and 32 missing observations; results for the remaining SBC conditions are shown in SOM Figures S27–29. Histograms show normalized posterior ranks of the true evolutionary VCV trace, intraspecific VCV traces, and taxon-mean elements. Under calibration, ranks are expected to be uniformly distributed. The annotation in the upper-right corner gives the smallest log(γ/γ_5_) among parameters in that category, followed by the number of parameters rejecting uniformity at the 5% level.

### Posterior estimates are well-sampled and data-consistent

We applied this model to two permanent teeth, the mandibular canine (C_1_) and maxillary second incisor (I^2^). For the C_1_ analysis with hominins (N_taxa_=8), we achieved a mean effective sample size (ESS) of 38,748 across all parameters within a single chain. The worst sampled parameter, the imputed decile 4 value for a *Po. pygmaeus* individual, had an ESS of 260. For the I^2^ analysis (N_taxa_=4), we achieved a mean ESS of 21,902 within a single chain. The worst sampled I^2^ parameter was the *Pan paniscus* decile 10 mean for which we achieved an ESS of 1,231. These high ESS values indicated that we had efficiently sampled from the posterior distribution in both analyses. We ensured convergence by calculating the rank-normalized Gelman-Rubin statistic (Vehtari et al., 2021) across four independent chains. The C_1_ analysis achieved an average 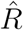 of 1.0003 and a maximum 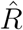 of 1.0048 across all parameters, and the I^2^ analysis achieved an average 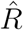 of 1.0002 and a maximum 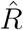 of 1.0008, well below the suggested threshold for convergence of 1.01 (Vehtari et al., 2021). All empirical results are reported from a single chain and are not pooled from these independent runs used to evaluate convergence. To ensure these posterior distributions are not prior-dominated, we calculated the Kullback-Leibler (KL) divergence between the prior and posterior distributions for our intraspecific and evolutionary VCV parameters (Kullback and Leibler, 1951). We see separation from the prior in all parameters, despite the small number of taxa and individuals per taxon, indicating that there is signal in our dataset about the model parameters despite these potential limitations (Table 1). The posterior distributions for the intraspecific VCV matrices inferred for *Pongo* and *Gorilla* exhibit the least separation from the prior inferred from the C_1_ dataset, and the *Pan paniscus* intraspecific VCV exhibits the least separation from the prior inferred from the I^2^ dataset (Table 1). This suggests that more individuals would help better characterize intraspecific variation in these taxa. Nevertheless, these diagnostics confirm that both analyses converged on their posterior distributions, efficiently sampled from those distributions, and that these posterior distributions are not prior-dominated, meaning our data are informative about the parameters of our model.

**Table 1.**
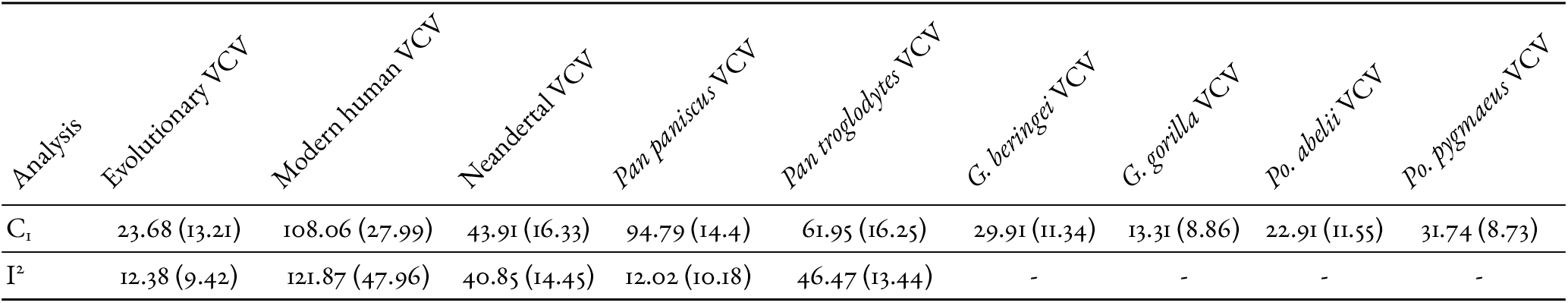
Kullback-Leibler divergences from the prior to the posterior distribution for the variance-covariance matrices. These posterior distributions are Inverse Wishart distributed; the maximum likelihood estimate for each posterior distribution’s degree of freedom is listed in parentheses. Lower degrees of freedom indicate heavier tails.

| Analysis | Evolutionary VCV | Modern human VCV | Neandertal VCV | *Pan paniscus* VCV | *Pan troglodytes* VCV | *G. beringei* VCV | *G. gorilla* VCV | *Po. abelii* VCV | *Po. pygmaeus* VCV |
| --- | --- | --- | --- | --- | --- | --- | --- | --- | --- |
| $C_i$ | 23.68 (13.21) | 108.06 (27.99) | 43.91 (16.33) | 94.79 (14.4) | 61.95 (16.25) | 29.91 (11.34) | 13.31 (8.86) | 22.91 (11.55) | 31.74 (8.73) |
| $I^2$ | 12.38 (9.42) | 121.87 (47.96) | 40.85 (14.45) | 12.02 (10.18) | 46.47 (13.44) | - | - | - | - |

We calculated the Fréchet variance for all *S* and Σ_evo_ parameters in both the C_1_ and I^2^ datasets. Fréchet variance measures uncertainty in these parameters while respecting the symmetric positive definite constraint on the matrices. For C_1_, the Fréchet variance in Σ_evo_ under the full model was 8.68, compared with 12.03 under the taxon means model; for I^2^, the corresponding values were 16.76 and 23.3 (Table 3). Among the intraspecific matrices, the modern human, *Pan paniscus*, and *Pan troglodytes* posterior distributions were the most dispersed in the C_1_ analysis, and the modern human and *Pan troglodytes* distributions were the most dispersed in the I^2^ analysis, with *Panpaniscus* the least. Because Σ_evo_ is estimated from only *m*−1 independent contrasts (7 for C_1_, 3 for I^2^) and shows among the lowest prior-to-posterior separation of any VCV parameter (Table 1), these evolutionary VCV posteriors should be read as informed but not sharply determined.

Finally, we employed posterior predictive checks to assess the fit of the model to our C_1_ and I^2^ datasets, and found that the model accurately reproduces the empirical structure of the dataset (SOM Figure S2-9). For the C_1_ analysis, 121 of 127 Bayesian *p*-values were within the acceptable range of between 0.05 and 0.95. We were unable to calculate a *p*-value for the *G. gorilla* D3 standard deviation as we only have one sample observed for that decile from that taxon. The Bayesian *p*-values for the *H. sapiens* D7-10 mean were all *<* 0.05, ranging between 0.0156 and 0.0404 (SOM Figure S2) indicating the posterior distribution supports a *H. sapiens* mean for those deciles lower than what is empirically observed. Additionally, the posterior distribution supports a larger standard deviation in *Pan paniscus* D3 and a smaller mean in *G. gorilla* D5 than what is empirically observed (SOM Figure S5-6). All Bayesian *p*-values were within 0.05 and 0.95 for the I^2^ analysis. The close correspondence between posterior predictive and empirical datasets suggests that the covariance structure is sufficiently identifiable from available data, and further suggests that the inferred covariance structures are not merely prior-regularized noise.

### Posterior estimates of intraspecific covariance differ among taxa

Under a model that estimates a distinct Σ_*i*_ for every taxon, we assessed the extent to which the resulting posterior distributions differed among taxa. Symmetrized KL divergence between Σ_*i*_ posterior distributions was ≥ 228.81 for every pair of congeneric taxa in both datasets, with the greatest divergences between modern humans and Neandertals (3,129.31 for C_1_; 3,765.93 for I^2^; Table 2). Symmetrized KL has no absolute scale, so we interpret these values comparatively: divergence between the two *Homo* taxa exceeds that between the two *Gorilla* taxa by more than an order of magnitude in C_1_, and within *Pan* the divergence is 1,206.84 for C_1_ but only 480.37 for I^2^. That ordering is consistent with the posterior predictive checks, which show the taxon-specific matrices reproducing per-taxon means and standard deviations and indicate that the perikymata data support unique intraspecific distributions for at least *Homo* and *Pan*.

**Table 2.** Symmetrized Kullback-Leibler divergences measuring the divergence in the inferred C_1_ and I^2^ intraspecific variance-covariance distributions between members of a genus.

| Comparison | Symmetrized KL divergence |
| --- | --- |
| $C_I$ | |
| Modern human vs. Neandertals | 3129.31 |
| *Pan paniscus* vs. *Pan troglodytes* | 1206.84 |
| *Gorilla beringei* vs. *Gorilla gorilla* | 228.81 |
| *Pongo abelii* vs. *Pongo pygmaeus* | 323.87 |
| $I^2$ | |
| Modern human vs. Neandertals | 3765.93 |
| *Pan paniscus* vs. *Pan troglodytes* | 480.37 |

To further compare our approach against inference with a single intraspecific VCV matrix, we calculated the geodesic distance on the symmetric positive definite manifold between each VCV matrix in the C_1_ posterior distribution and its corresponding maximum likelihood estimate under the Felsenstein (2008) model as implemented in Goolsby et al. (2017) (SOM Figure S1). No matrix in our C_1_ posterior distribution was similar to the evolutionary VCV or the singular intraspecific VCV estimated under the Felsenstein (2008) model; in each case the maximum likelihood estimate lay farther from the posterior Fréchet mean than every single posterior sample (SOM Figure S1). The greatest distances were between the evolutionary VCV posterior and the Felsenstein (2008) maximum-likelihood evolutionary VCV.

### Modeling taxon-specific intraspecific structure changes the inferred evolutionary process

By fitting a multivariate Brownian motion to the estimated mean PMD, we modeled the evolution of the mean PMD for each taxon on the great ape phylogenetic tree (Figure 3). Jointly inferring this multivariate Brownian motion alongside taxon-specific intraspecific VCV matrices yields mean evolutionary VCV matrices that differ substantially, for both the C_1_ and the I^2^, from those inferred with species means fixed.

**Figure 3:**
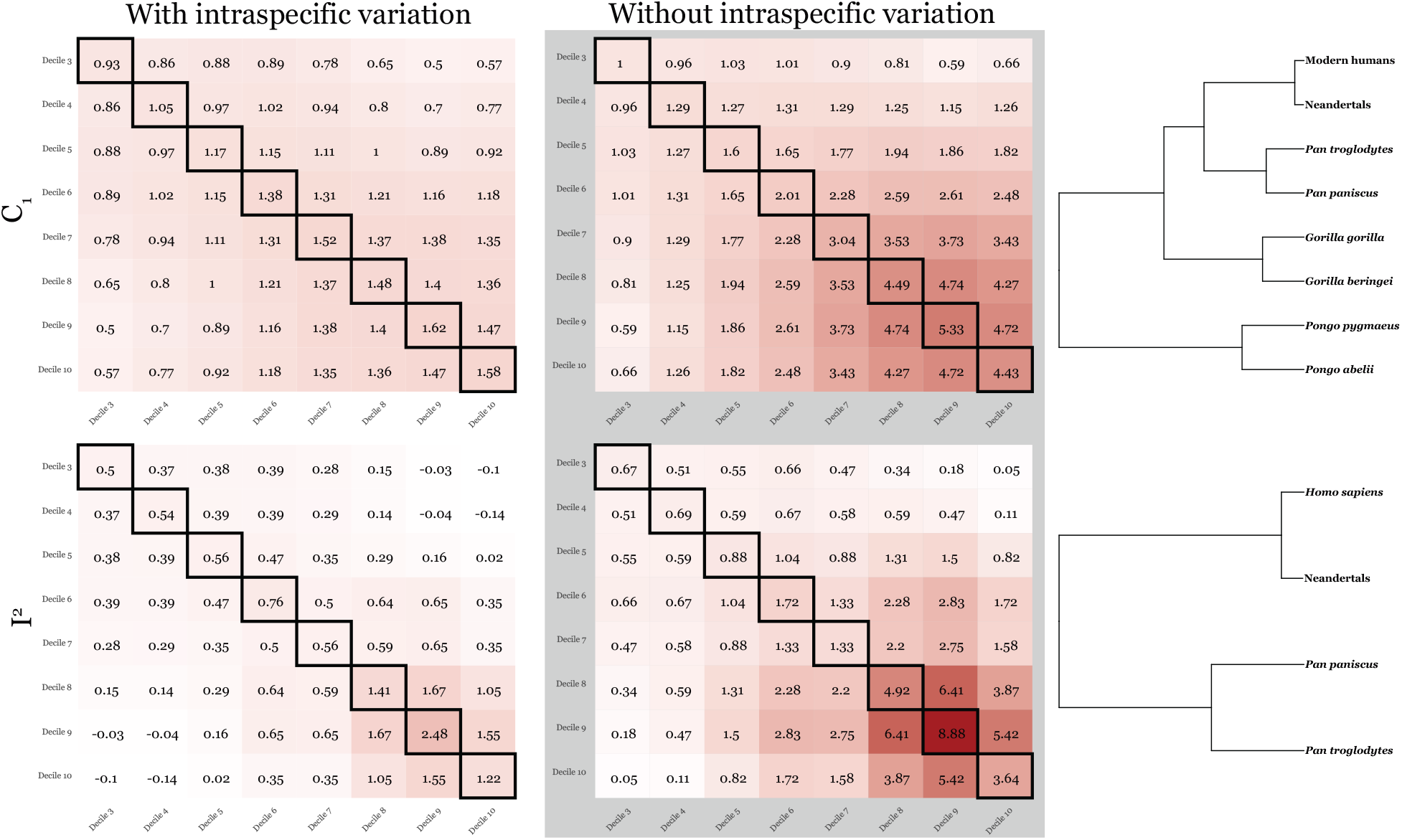
Posterior mean evolutionary variance-covariance matrices. Top row, results from the C_1_ analysis; bottom row, results from the I^2^ analysis. Left shows the mean evolutionary variance-covariance matrix that describes the evolution of the intraspecific means on the corresponding great ape phylogeny (right). Center shows the mean evolutionary variance-covariance matrix estimated without intraspecific variation from fixed species means. Diagonal elements indicate the evolutionary rate of change for that decile and are bordered with black. Off diagonal elements indicate the evolutionary covariance across deciles. Darker colors indicate higher rates or covariances. Each mean variance-covariance matrix was calculated on the SPD manifold.

For both teeth, the evolutionary VCV matrices inferred without intraspecific variation show cervical deciles (D6-10) evolving considerably faster than cuspal deciles (D3-5), and show positive evolutionary covariances for every pair of deciles (Figure 3). When the same matrices are inferred alongside taxon-specific intraspecific distributions, both features change. Evolutionary rates become far more uniform across the crown, ranging from 0.93 to 1.62 in the C_1_ and 0.5 to 2.48 in the I^2^, compared with 1 to 5.33 and 0.67 to 8.88 respectively under the taxon means model (Figure 3). Although the prior expectation for each rate is 1, the inferred off-diagonal elements depart markedly from their prior expectation of ≈ 0, reaching 1.47 in the C_1_ and 1.67 in the I^2^ (Figure 3); the shift toward more uniform rates is therefore not simply regression of the posterior toward the prior mean.

Notably, the deciles whose rates are largest under the taxon means model are also among those with the greatest intraspecific variance (Table 4), consistent with variation that is not attributed to within-taxon uncertainty being instead attributed to the evolutionary component. The evolutionary covariances change more dramatically still in the I^2^. Under the full model, the two most cuspal deciles retained (D3-4) show negative inferred evolutionary covariances with the late cervical deciles (D9-10), reaching −0.14 (D4 and D10), alongside strong positive covariance within the late cervical deciles (1.67 between D8 and D9, 1.55 between D9 and D10; Figure 3). Every element of the corresponding taxon means matrix is positive. Fixing species means increased rate estimates for both the C_1_ and I^2^ and, in the I^2^, reversed the sign of the inferred evolutionary relationship between the most cuspal and the late cervical deciles.

These differences are not artifacts of summarizing each posterior by its mean. Comparing the element-wise marginal posterior distributions between the two models, rates and covariances corresponding to cuspal and mid-crown deciles show the highest overlap, while those for cervical deciles show the least (SOM Table S2). For the C_1_, the marginal posterior distribution for the rate of D3 overlaps 95.8% between the full and taxon means models, while the rate for D9 overlaps 40.59%. The same gradient is present, though weaker, in the I^2^, where the corresponding overlaps are 94.37% (D4) and 68.82% (D9), consistent with the smaller number of independent contrasts available for that tooth. We therefore inferred distinct evolutionary VCV posterior distributions between the two models for both tooth types, not merely distinct *post hoc* summaries of those distributions.

### Comparisons of intraspecific distributions among taxa

We investigated enamel growth differences between modern humans and Neandertals through samples from the posterior predictive distribution, which integrates over uncertainty in both the taxon-specific mean and the taxon-specific variance-covariance matrix. The posterior predictive distribution is the probability distribution over unobserved samples, conditioned on the observed samples and the inferred parameters of our model. It is therefore the probability distribution over all Neandertals, modern humans, and great ape individuals not included in our dataset. This posterior predictive distribution accounts for all the sources of uncertainty in our data and provides informed estimates of overall population-wide PMD in each taxon included here.

Modern human and Neandertal posterior predictive PMD (ppPMD) for the I^2^ show a pattern of high overlap (≥ 66.1%) across D3-7, with modern humans and Neandertals initially separating at D8 (46.6%), and remaining separated through D9 (40.2%) and D10 (37.1%) (Figure 4). In contrast, for C_1_ there is higher overlap of ppPMD (≥ 68.3%) in cuspal deciles (D3-5) (Figure 4). They begin to differentiate at D6, with an overlap of 53.9%, and further diverge in D7-9, reaching a minimum overlap of 28.5% at D8 (Figure 4). However, in D10, modern human and Neandertal ppPMD reverse this pattern with an overlap of 57.1%. This reversal is driven primarily by increasing ppPMD variance in both taxa at D10 rather than by convergence of their means (Figure 4, Table 4). Thus, it is within D7-9 that modern human and Neandertal canine ppPMD are most distinguishable.

**Figure 4:**
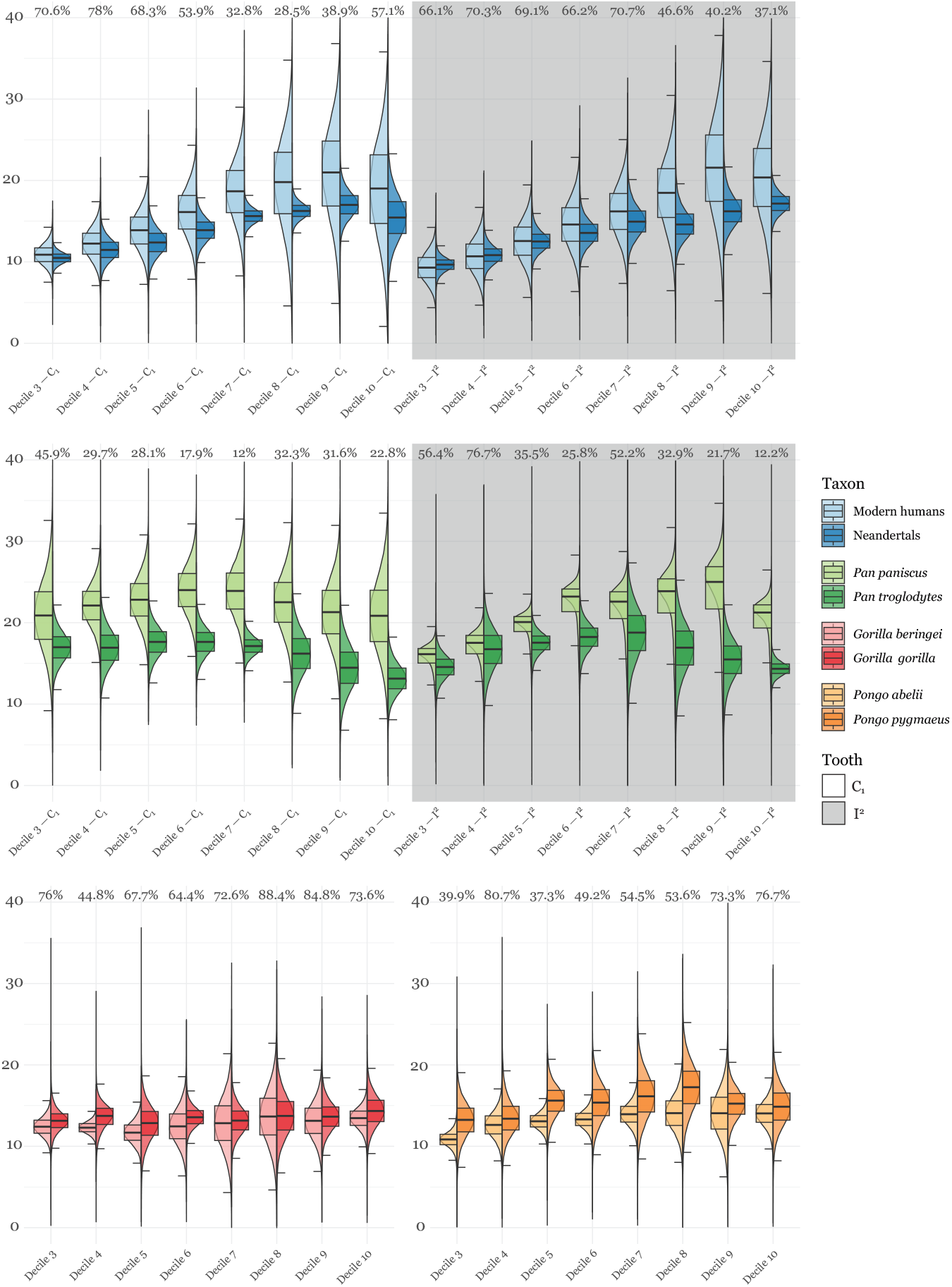
Draws from the posterior predictive distribution for each taxon compared. Within each genus examined—*Homo, Pan, Gorilla*, and *Pongo*— we directly compare the posterior predictive distribution describing the C_1_ (white background) and I^2^ (gray background) perikymata per millimeter per decile between the two groups we sampled for each genus. Additionally within each genus for each decile, we calculate the percent of the posterior predictive distribution overlapping between each group within the genus. Overlap percentage is calculated as the area occupied by the minimum of the two posterior predictive distributions. All posterior predictive distributions are plotted truncated at zero and forty for visual clarity. The boxplot center line corresponds to the median, box limits the upper and lower quartiles, whiskers the maximum and minimum posterior predictive sample within 1.5 *×* the inter-quartile range.

Non-human great apes present a varied picture with respect to their I^2^ and C_1_ development. The *Pan troglodytes* and *Pan paniscus* I^2^ ppPMD range from remarkably similar (76.7% for D4) to substantially separated for D9 and D10 (21.7% and 12.2% overlap, respectively), though we note that the I^2^ *Panpaniscus* sample comprises only two individuals. The two species diverge consistently in ppPMD for the C_1_, with overlaps between 12% (D7) and 45.9% (D3) (Figure 4). In sharp contrast to *Pan*, taxa within *Gorilla* and *Pongo* exhibited some of the greatest ppPMD overlaps (Figure 4). *G. beringei* and *G. gorilla* C_1_ overlap at least 64.4% in all deciles except D4 (44.8%). In *Pongo, Po. abelii* and *Po. pygmaeus* C_1_ ppPMD overlaps range from 37.3% overlap in D5 to 80.7% in D4.

## Discussion

Here, we introduced a hierarchical Bayesian framework that jointly infers the evolutionary covariance structure alongside a separate intraspecific covariance structure for every taxon. We confirmed that both the model and its implementation in BURL yield well-calibrated posterior distributions across 144 combinations of tree size, trait dimensionality, sample size, and missingness, including regimes in which traits outnumber individuals per taxon (a known inferential challenge; Clavel et al., 2019). Prior-to-posterior separation in every VCV matrix inferred in our empirical application indicates that these estimates are driven by the data rather than the prior, even where the matrices are severely under-determined. Through several lines of evidence, we confirmed that our framework explains the observed structure of perikymata spacing within each great ape taxon and that the inferred intraspecific covariance structures differ among congeneric taxa to a degree that a single shared matrix could not represent. The calibration and empirical results connect directly: because the fixed-means procedure was systematically under-calibrated under the generative conditions examined here, empirical differences between the two approaches should not be interpreted without accounting for uncertainty in taxon means. The substantial differences we observe between the two inferred evolutionary VCV matrices are consistent with a redistribution of inferred variation from the evolutionary to the intraspecific component once uncertainty in taxon means is modeled. This correction is one that BURL can make in a dataset as difficult to sample as great ape PMD, where several taxa are represented by fewer individuals than traits and missing data are common.

### Methodology

The robustness in our method across both the simulated and empirical datasets is enabled through our application of the inverse Wishart conjugate relationship to the multivariate normal distribution, which allows us to directly sample from the posterior distribution for these parameters with corresponding computational benefits (Gelman et al., 2013). We also exploit this conjugate relationship to aid inference of the evolutionary VCV matrix describing the multivariate Brownian motion process. The pairing of multivariate Brownian motion and the inverse Wishart process was likewise made due to its computational advantages. Alternative inferential strategies, such as a separation strategy decomposing each VCV matrix into a vector of variances and a correlation matrix also exist, but introduce further inferential complexity that our Gibbs sampler avoids (Caetano and Harmon, 2019; Fisher et al., 2021). Our choice of the inverse Wishart prior reflects a deliberate trade-off between modeling flexibility and computational efficiency.

This approach of modeling a variance parameter at each taxon can be seen as an extension of Kostikova et al. (2016) to multivariate continuous traits, though unlike Kostikova et al. (2016), we do not consider the evolution of variance itself. We were able to relax the assumptions of prior multivariate phylogenetic comparative models (e.g., Adams and Collyer, 2024; Felsenstein, 2008) and explicitly model unique intraspecific distributions for each tip. Our observation of substantial differences in inferred intraspecific VCV matrices, and within the *Homo* and *Pan* genera in particular, reinforces the need to relax these assumptions.

## Empirical results

Our empirical data provides real world support that our hierarchical model can appropriately partition variation between intraspecific and interspecific processes. We restricted our simulation study design to compare our full model against the taxon means model as, to our knowledge, there is not a Bayesian implementation of the Felsenstein (2008) model; as such, a direct, coverage-based comparison between our full model and a single, shared intraspecific VCV matrix is not straightforward. However, by utilizing geodesic distances on the SPD manifold, we were able to compare inferred VCV matrices from our full model posterior distribution against maximum-likelihood point estimates under the Felsenstein (2008) (as implemented in Goolsby et al. 2017) while respecting their underlying mathematical structure. These analyses demonstrate that allowing taxon-specific covariance structure yields evolutionary and intraspecific covariance estimates that differ substantially from those obtained under fixed-means or shared-covariance procedures. They do not, however, constitute formal model comparison; marginal likelihoods, cross-validation, or another predictive model-comparison framework would be required to establish relative model support.

Fréchet variances have proven to be a useful tool to describe the uncertainty in the posterior distributions for *S*, Σ_evo_ in a way that respects the symmetric positive definite structure of these matrices. Together, the Fréchet variances and prior-to-posterior KL divergences jointly characterize both the information available for each VCV matrix and the residual uncertainty in it, and three patterns emerge in the C_1_ dataset. First, modern humans and *Pan paniscus* have both the highest KL divergences and the highest Fréchet variances of any parameter (Table 1, Table 3). This pairing indicates that not only are the data highly informative towards the intraspecific VCV matrices for these parameters, but that our posterior distributions may be recovering genuine biological variability in the VCV matrices that drives the diffuse posterior distributions. Second, Neandertal and *Pan troglodytes* both exhibit relatively high KL divergences and moderate Fréchet variances, indicating informed inference of potentially less variable VCV matrices compared to the hominins. Finally, *Gorilla* and *Pongo* both exhibit relatively low KL divergences and low to moderate Fréchet variances. However, in this case, the latter may result from the influence of the prior on the posterior estimation, particularly in *G. gorilla*, the taxon with the lowest KL divergence in the intraspecific VCV. Similarly, in the I^2^ analysis, the posterior distribution is most uncertain for the modern human and *Pan troglodytes* intraspecific VCV matrices; these taxa, particularly modern humans, have the highest KL divergences (Table 1, Table 3). Likewise, both the Fréchet variance and KL divergence for the *Pan paniscus* intraspecific VCV are the lowest among all I^2^ intraspecific VCV matrices, suggesting that more data would better clarify whether this lack of variance is truly indicative of a lack of biological variability (Table 1, Table 3).

**Table 3.** Fréchet variance for each VCV posterior distribution. The evolutionary VCV is compared between the full model, which accounts for intraspecific variation, and the taxon means model, in which species means are fixed at their observed sample means and only the evolutionary VCV is estimated; the remaining columns give the intraspecific VCV inferred for each taxon under the full model, which has no analog under the taxon means model. Higher values indicate greater uncertainty in the inferred VCV matrix.

| Analysis | Evolutionary VCV |  | Intraspecific VCV (full model) |  |  |  |  |  |  |  |
| --- | --- | --- | --- | --- | --- | --- | --- | --- | --- | --- |
|  | Full model | Taxon means model | Modern human | Neandertal | *Pan paniscus* | *Pan troglodytes* | *G. beringei* | *G. gorilla* | *Po. abelii* | *Po. pygmaeus* |
| $C_1$ | 8.68 | 12.03 | 78.84 | 44.66 | 90.35 | 58.68 | 38.31 | 26.86 | 26.24 | 45.33 |
| $I^2$ | 16.76 | 23.3 | 75.82 | 35.85 | 22.47 | 62.97 | - | - | - | - |

Estimating a full intraspecific distribution for each taxon lets us ask not only whether taxa differ in mean perikymata spacing, but whether they differ in how variable that spacing is. Intraspecific variation in modern humans and Neandertals is consistently divergent between the two taxa across both the C_1_ and I^2^; much of the overlap between the two taxa in their later-forming deciles of both teeth is likely due to the substantial and increasing variance in the modern human ppPMD for those deciles. Neandertals have some of the lowest variances in both tooth types of any taxon except for D10 in their canine, where variance increases 3-fold relative to D9. Thus, while the ppPMD means for modern humans and Neandertals remain distant for these later-forming deciles, it is the increase in variance in both taxa that is driving the increased overlap in the C_1_ D10 (Figure 4, Table 4). We also note that the modern human C_1_ posterior predictive means for D7-10 fall below their empirical counterparts (SOM Figure S2). As modern humans are the most extreme taxon in this clade for cervical PMD, we speculate that this reflects modest shrinkage of the inferred mean toward the remainder of the tree under the Brownian prior on *M*; if so, the modern human–Neandertal separation we report for these deciles is, if anything, conservative. Notably, modern human and Neandertal intraspecific VCV posterior distributions are more divergent than any other pair of taxa across both tooth types considered (Table 2), consistent with distinct inferred patterns of intraspecific covariance.

**Table 4.**
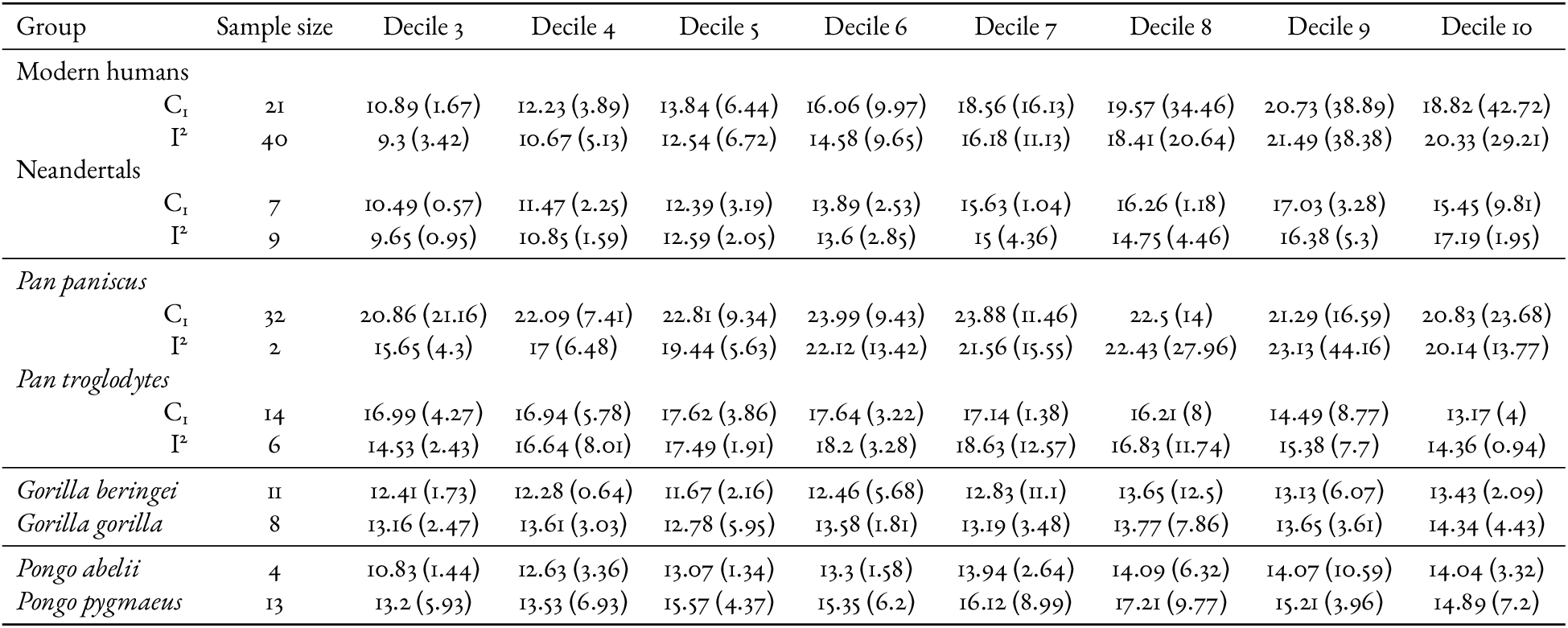
Comparison of posterior predictive perikymata per millimeter per decile means and variances (in parentheses). For modern humans, Neandertals, and *Pan*, where we have both C_1_ and I^2^ data, both values are listed. We only have C_1_ data for *Gorilla* and *Pongo*.

| Group | Sample size | Decile 3 | Decile 4 | Decile 5 | Decile 6 | Decile 7 | Decile 8 | Decile 9 | Decile 10 |
| --- | --- | --- | --- | --- | --- | --- | --- | --- | --- |
| Modern humans |  |  |  |  |  |  |  |  |  |
| C<sub>1</sub> | 21 | 10.89 (1.67) | 12.23 (3.89) | 13.84 (6.44) | 16.06 (9.97) | 18.56 (16.13) | 19.57 (34.46) | 20.73 (38.89) | 18.82 (42.72) |
| I<sup>2</sup> | 40 | 9.3 (3.42) | 10.67 (5.13) | 12.54 (6.72) | 14.58 (9.65) | 16.18 (11.13) | 18.41 (20.64) | 21.49 (38.38) | 20.33 (29.21) |
| Neandertals |  |  |  |  |  |  |  |  |  |
| C<sub>1</sub> | 7 | 10.49 (0.57) | 11.47 (2.25) | 12.39 (3.19) | 13.89 (2.53) | 15.63 (1.04) | 16.26 (1.18) | 17.03 (3.28) | 15.45 (9.81) |
| I<sup>2</sup> | 9 | 9.65 (0.95) | 10.85 (1.59) | 12.59 (2.05) | 13.6 (2.85) | 15 (4.36) | 14.75 (4.46) | 16.38 (5.3) | 17.19 (1.95) |
| *Pan paniscus* |  |  |  |  |  |  |  |  |  |
| C<sub>1</sub> | 32 | 20.86 (21.16) | 22.09 (7.41) | 22.81 (9.34) | 23.99 (9.43) | 23.88 (11.46) | 22.5 (14) | 21.29 (16.59) | 20.83 (23.68) |
| I<sup>2</sup> | 2 | 15.65 (4.3) | 17 (6.48) | 19.44 (5.63) | 22.12 (13.42) | 21.56 (15.55) | 22.43 (27.96) | 23.13 (44.16) | 20.14 (13.77) |
| *Pan troglodytes* |  |  |  |  |  |  |  |  |  |
| C<sub>1</sub> | 14 | 16.99 (4.27) | 16.94 (5.78) | 17.62 (3.86) | 17.64 (3.22) | 17.14 (1.38) | 16.21 (8) | 14.49 (8.77) | 13.17 (4) |
| I<sup>2</sup> | 6 | 14.53 (2.43) | 16.64 (8.01) | 17.49 (1.91) | 18.2 (3.28) | 18.63 (12.57) | 16.83 (11.74) | 15.38 (7.7) | 14.36 (0.94) |
| *Gorilla beringei* | 11 | 12.41 (1.73) | 12.28 (0.64) | 11.67 (2.16) | 12.46 (5.68) | 12.83 (11.1) | 13.65 (12.5) | 13.13 (6.07) | 13.43 (2.09) |
| *Gorilla gorilla* | 8 | 13.16 (2.47) | 13.61 (3.03) | 12.78 (5.95) | 13.58 (1.81) | 13.19 (3.48) | 13.77 (7.86) | 13.65 (3.61) | 14.34 (4.43) |
| *Pongo abelii* | 4 | 10.83 (1.44) | 12.63 (3.36) | 13.07 (1.34) | 13.3 (1.58) | 13.94 (2.64) | 14.09 (6.32) | 14.07 (10.59) | 14.04 (3.32) |
| *Pongo pygmaeus* | 13 | 13.2 (5.93) | 13.53 (6.93) | 15.57 (4.37) | 15.35 (6.2) | 16.12 (8.99) | 17.21 (9.77) | 15.21 (3.96) | 14.89 (7.2) |

In contrast to the pattern observed in modern humans and Neandertals, the relationship between *Pan* species is strikingly different between the I^2^ and C_1_. *Pan troglodytes* and *Pan paniscus* diverge in C_1_ PMD, which appears to indicate that we are recovering a basic difference in enamel deposition between the two species of *Pan*. However, the broadly declining (though not monotonically so) overlap between *Pantroglodytes* and *P. paniscus* observed in I^2^ is shared. Canines are more sexually dimorphic than incisors and so the variation we identify could be due to an imbalance in sex sampling between the two species or reflect different developmental rates in the late juvenile period between the species regardless of sex.

The overlap in C_1_ PMD between the two *Gorilla* and the two *Pongo* species add to this complicated picture. In neither genus is there any evidence for divergences across multiple deciles of the broad, systematic scale we observed in *Homo* and *Pan*. Further suggesting extensive conservatism within *Pongo* and *Gorilla* are the symmetrized KL divergence results, which demonstrate greater similarity in the posterior distributions within those two genera compared to *Homo* and *Pan* (Table 2). However, we do note that KL divergences comparing posterior to prior distribution are the lowest in the intraspecific VCV matrices inferred for members of *Pongo* and *Gorilla*, suggesting limited information in the data towards these parameters (Table 1). Thus, while our results suggest conservation in the pattern of enamel deposition in *Pongo* and *Gorilla*, more data would increase confidence.

### Limitations

Several aspects of our data and our modeling choices constrain inferences from these. Perikymata are natively an ordinal, integer manifestation (i.e., a count) of several underlying continuous processes, making standard phylogenetic models difficult to apply. We circumvented this limitation by scaling perikymata count per decile by the height of that decile. However, while the resulting PMD is a continuous trait, it is only biologically meaningful when PMD is positive. This biological bound created a model misspecification between our data and our models, as multivariate Brownian motion and multivariate normal distributions have an unbounded support and therefore assign positive (albeit low) probability to biologically unrealistic species means, histories of change that include unrealistic species means, and biologically unrealistic imputed missing PMD. Unfortunately, applying truncated multivariate normal distributions to a phylogeny (and by extension, truncated multivariate Brownian motion processes) is nontrivial (Botev, 2017; Damien and Walker, 2001), hindering our ability to match the biological bounds in our inference. Future work may clarify the impact of these bounding decisions. Given that our intraspecific mean posterior distributions and posterior predictive distributions were centered well above zero, the probability mass assigned below zero was small and likely created a negligible bias in our results.

Additionally, we only assessed multivariate Brownian motion; alternative models like Ornstein-Uhlenbeck models (Hansen, 1997) may better explain the distribution of perikymata across the great ape clade. We did not perform a model fitting analysis (e.g., Lartillot and Philippe, 2006; Xie et al., 2011) as the goal of this study was not to identify the best-fit model to perikymata, but instead describe a framework for tractable joint estimation of evolutionary process and intraspecific distributions in multivariate data. Additionally, these alternative models introduce computational complexity, as there is not a clear conjugate relationship that we could exploit. Thus, in this inferential regime of few taxa where model-fitting is already badly underpowered in the simplistic multivariate Brownian motion case, more complex models like the Ornstein-Uhlenbeck would be further underpowered regardless of implementation. Future work may relax this multivariate Brownian motion assumption; importantly, this hierarchical framework is independent of the evolutionary model, thus making this future work easy to implement.

Finally, each Σ_*i*_ receives an independent inverse Wishart prior, so closely related taxa are no more similar in covariance structure *a priori* than distantly related ones. Phylogenetic models describing the evolution of variance-covariance matrices exist, but are computationally expensive with limited inferential power (e.g., Blomberg et al., 2025), which currently prevents a full extension of Kostikova et al. (2016) to multivariate inference. Extending the hierarchy so that intraspecific covariance structure itself evolves along the phylogeny is nonetheless the natural next step, and one our framework’s separation of the evolutionary and intraspecific levels would accommodate directly.

## Conclusions

The hierarchical framework we present here represents a step toward fully integrating inter- and intraspecific variation within a unified comparative analysis. In particular, this model provides a principled way to estimate species means as full posterior distributions, rather than as point estimates. Using species means as fixed and known is common practice in phylogenetic comparative methods (Garamszegi and Møller, 2010), but our simulations demonstrate that treating species means as fixed can be overconfident and yield biased uncertainty estimates under the conditions examined here. Our model’s ability to estimate true species means given uncertainty both in evolutionary process and intraspecific variation is especially consequential in datasets like ours, where intraspecific sample sizes are small and missing data is commonplace, a scenario particularly common in paleontological contexts. We hope this framework makes propagating intraspecific uncertainty through comparative analyses standard practice rather than the exception, and that it brings phylogenetic and population-level inference one step closer together.

## Data availability

The code needed to replicate the findings of this study is openly available on GitHub at https://github.com/Levi-Raskin/PerikymataPhylogenetics. This repository includes the version of BURL used for all analyses and all associated scripts needed for subsequent analysis. The data that support the findings of this study are openly available on Dryad at http://doi.org/10.5061/dryad.nzs7h4565.

## Funding

This material is based upon work supported by the U.S. Department of Energy, Office of Science, Office of Advanced Scientific Computing Research, under Award Number DE-SC0026073 to LYR. This report was prepared as an account of work sponsored by an agency of the United States Government. Neither the United States Government nor any agency thereof, nor any of their employees, makes any warranty, express or implied, or assumes any legal liability or responsibility for the accuracy, completeness, or usefulness of any information, apparatus, product, or process disclosed, or represents that its use would not infringe privately owned rights. Reference herein to any specific commercial product, process, or service by trade name, trademark, manufacturer, or otherwise does not necessarily constitute or imply its endorsement, recommendation, or favoring by the United States Government or any agency thereof. The views and opinions of authors expressed herein do not necessarily state or reflect those of the United States Government or any agency thereof.

## Acknowledgments

We are grateful to Drs. Rasmus Nielsen, Jack Tseng, and Narimane Chatar for their helpful comments and feedback during the preparation of this manuscript. We are grateful to Dr. Donald Reid for sharing data. We thank Drs. Kristin Krueger, Adam Ferguson, and Larry Heaney for their assistance molding and casting. We also thank Dr. Zak Kerrigan for his microscopy support.

## Notes

https://github.com/Levi-Raskin/PerikymataPhylogenetics

http://doi.org/10.5061/dryad.nzs7h4565

### Competing Interest Statement

The authors have declared no competing interest.

### Summary of Updates

Added simulation based calibration results to the main text of the manuscript and restructured the manuscript to highlight the methodological contribution more.

