## Supplementary Information for "A hierarchical Bayesian framework accommodates intraspecific and interspecific variation in multivariate traits"

Levi Yoder Raskin[a,b,c,1], Maja Šešelj[b], John Huelsenbeck[a], Wonseop Lim[a], Jacky Kaiyuan Li[d], Debbie Guatelli-Steinberg[e], Mackie C. O'Hara[f,2], and Bárbara Domingues Bitarello[c,2]

[a]Department of Integrative Biology, University of California, Berkeley

[b]Department of Anthropology, Bryn Mawr College

[c]Department of Biology, Bryn Mawr College

[d]Biostatistics Division, University of California, Berkeley

[e]Department of Anthropology, The Ohio State University

[f]Department of Biology, Ball State University

September 2, 2026

**Table 1:** Coverage probability and Markov chain Monte Carlo (MCMC) runtime from a full-factorial simulation study that varied the number of taxa (8, 16, or 32; shown top to bottom in separate panels), the number of traits (8, 16, or 32; panel rows), the number of individuals per taxon (2, 4, 8, or 16; panel columns), and the number of missing observations across the entire simulated dataset (0, 8, 16, or 32; rows within each cell). For each combination of conditions we simulated 100 datasets and inferred parameters with both the full model, which accounts for intraspecific variation, and the taxon means model, which takes the mean value across individuals in a taxon as known without uncertainty. Within each cell, the top line reports coverage probability (mean ± standard error across replicates) for the full model (left of the slash) and the taxon means model (right of the slash); the bottom line reports the mean wall clock time in minutes per ten million MCMC cycles for the full model running serially. The expected coverage probability is 95%.

| | | Individuals per Taxon | | | |
|---|---|---|---|---|---|
| **Traits** | **Missing** | **2** | **4** | **8** | **16** |
| | | **8 Taxa** | | | |
| 8 | 0 | $0.95 \pm 0.004 / 0.64 \pm 0.019$<br>0.8 m | $0.96 \pm 0.003 / 0.72 \pm 0.016$<br>0.9 m | $0.95 \pm 0.003 / 0.78 \pm 0.016$<br>1.0 m | $0.95 \pm 0.003 / 0.84 \pm 0.012$<br>1.2 m |
| 8 | 8 | $0.95 \pm 0.003 / 0.65 \pm 0.017$<br>0.8 m | $0.95 \pm 0.003 / 0.71 \pm 0.015$<br>0.8 m | $0.95 \pm 0.003 / 0.75 \pm 0.014$<br>0.9 m | $0.94 \pm 0.003 / 0.83 \pm 0.014$<br>1.2 m |
| 8 | 16 | $0.95 \pm 0.004 / 0.65 \pm 0.017$<br>0.8 m | $0.95 \pm 0.003 / 0.72 \pm 0.015$<br>0.8 m | $0.95 \pm 0.002 / 0.77 \pm 0.014$<br>0.9 m | $0.95 \pm 0.003 / 0.82 \pm 0.017$<br>1.2 m |
| 8 | 32 | $0.95 \pm 0.003 / 0.64 \pm 0.015$<br>0.8 m | $0.94 \pm 0.003 / 0.72 \pm 0.014$<br>0.8 m | $0.95 \pm 0.003 / 0.77 \pm 0.016$<br>1.0 m | $0.95 \pm 0.002 / 0.81 \pm 0.015$<br>1.2 m |
| 16 | 0 | $0.95 \pm 0.003 / 0.69 \pm 0.014$<br>2.2 m | $0.95 \pm 0.003 / 0.74 \pm 0.012$<br>2.4 m | $0.95 \pm 0.003 / 0.79 \pm 0.015$<br>2.8 m | $0.95 \pm 0.003 / 0.83 \pm 0.015$<br>3.3 m |
| 16 | 8 | $0.95 \pm 0.003 / 0.70 \pm 0.015$<br>2.2 m | $0.95 \pm 0.003 / 0.72 \pm 0.016$<br>2.5 m | $0.95 \pm 0.003 / 0.77 \pm 0.018$<br>2.7 m | $0.95 \pm 0.003 / 0.85 \pm 0.015$<br>3.3 m |
| 16 | 16 | $0.95 \pm 0.003 / 0.67 \pm 0.015$<br>2.3 m | $0.95 \pm 0.003 / 0.74 \pm 0.012$<br>2.4 m | $0.95 \pm 0.003 / 0.79 \pm 0.012$<br>2.7 m | $0.95 \pm 0.002 / 0.85 \pm 0.011$<br>3.3 m |

*continued on next page*

*Table 1 continued from previous page*

| | | Individuals per Taxon | | | |
|---|---|---|---|---|---|
| **Traits** | **Missing** | **2** | **4** | **8** | **16** |
| 16 | 32 | 0.95 ± 0.003 / 0.70 ± 0.013<br>2.3 m | 0.95 ± 0.003 / 0.74 ± 0.015<br>2.5 m | 0.95 ± 0.003 / 0.80 ± 0.013<br>2.7 m | 0.95 ± 0.002 / 0.84 ± 0.011<br>3.3 m |
| 32 | 0 | 0.95 ± 0.003 / 0.69 ± 0.015<br>11.0 m | 0.95 ± 0.003 / 0.75 ± 0.013<br>11.5 m | 0.95 ± 0.003 / 0.80 ± 0.015<br>12.5 m | 0.95 ± 0.003 / 0.87 ± 0.011<br>14.2 m |
| 32 | 8 | 0.95 ± 0.004 / 0.72 ± 0.010<br>11.1 m | 0.95 ± 0.003 / 0.76 ± 0.013<br>11.5 m | 0.95 ± 0.003 / 0.80 ± 0.013<br>12.5 m | 0.95 ± 0.003 / 0.85 ± 0.010<br>14.2 m |
| 32 | 16 | 0.95 ± 0.004 / 0.68 ± 0.015<br>10.9 m | 0.95 ± 0.003 / 0.75 ± 0.012<br>11.6 m | 0.95 ± 0.002 / 0.82 ± 0.010<br>12.2 m | 0.95 ± 0.003 / 0.86 ± 0.012<br>14.1 m |
| 32 | 32 | 0.95 ± 0.003 / 0.68 ± 0.012<br>11.1 m | 0.94 ± 0.003 / 0.73 ± 0.012<br>11.4 m | 0.95 ± 0.003 / 0.80 ± 0.014<br>12.1 m | 0.95 ± 0.003 / 0.84 ± 0.011<br>14.1 m |
| | | | **16 Taxa** | | |
| 8 | 0 | 0.95 ± 0.003 / 0.65 ± 0.015<br>0.9 m | 0.95 ± 0.003 / 0.68 ± 0.016<br>0.9 m | 0.95 ± 0.002 / 0.73 ± 0.015<br>1.1 m | 0.95 ± 0.002 / 0.78 ± 0.017<br>1.4 m |
| 8 | 8 | 0.95 ± 0.002 / 0.66 ± 0.016<br>0.9 m | 0.95 ± 0.002 / 0.70 ± 0.013<br>1.0 m | 0.95 ± 0.002 / 0.76 ± 0.014<br>1.1 m | 0.95 ± 0.002 / 0.82 ± 0.013<br>1.4 m |
| 8 | 16 | 0.95 ± 0.002 / 0.64 ± 0.014<br>0.9 m | 0.95 ± 0.002 / 0.69 ± 0.017<br>0.9 m | 0.95 ± 0.002 / 0.71 ± 0.017<br>1.1 m | 0.95 ± 0.002 / 0.79 ± 0.016<br>1.4 m |
| 8 | 32 | 0.95 ± 0.003 / 0.65 ± 0.015<br>0.9 m | 0.95 ± 0.002 / 0.67 ± 0.017<br>0.9 m | 0.94 ± 0.003 / 0.74 ± 0.014<br>1.1 m | 0.95 ± 0.002 / 0.78 ± 0.013<br>1.3 m |
| 16 | 0 | 0.95 ± 0.002 / 0.67 ± 0.015<br>2.5 m | 0.94 ± 0.002 / 0.74 ± 0.011<br>2.8 m | 0.95 ± 0.002 / 0.75 ± 0.016<br>3.0 m | 0.95 ± 0.002 / 0.82 ± 0.011<br>3.6 m |

*continued on next page*

*Table 1 continued from previous page*

| | | Individuals per Taxon | | | |
|---|---|---|---|---|---|
| **Traits** | **Missing** | **2** | **4** | **8** | **16** |
| 16 | 8 | $0.95 \pm 0.002$ / $0.70 \pm 0.012$<br>2.5 m | $0.95 \pm 0.002$ / $0.70 \pm 0.017$<br>2.8 m | $0.95 \pm 0.002$ / $0.77 \pm 0.014$<br>3.0 m | $0.95 \pm 0.002$ / $0.83 \pm 0.012$<br>3.6 m |
| 16 | 16 | $0.95 \pm 0.002$ / $0.70 \pm 0.013$<br>2.6 m | $0.95 \pm 0.002$ / $0.71 \pm 0.015$<br>2.7 m | $0.95 \pm 0.002$ / $0.78 \pm 0.012$<br>3.0 m | $0.95 \pm 0.002$ / $0.83 \pm 0.011$<br>3.6 m |
| 16 | 32 | $0.95 \pm 0.002$ / $0.70 \pm 0.013$<br>2.6 m | $0.95 \pm 0.002$ / $0.74 \pm 0.012$<br>2.8 m | $0.95 \pm 0.002$ / $0.77 \pm 0.012$<br>3.1 m | $0.95 \pm 0.002$ / $0.81 \pm 0.012$<br>3.6 m |
| 32 | 0 | $0.95 \pm 0.002$ / $0.71 \pm 0.013$<br>12.9 m | $0.95 \pm 0.002$ / $0.73 \pm 0.012$<br>13.1 m | $0.95 \pm 0.002$ / $0.78 \pm 0.015$<br>14.0 m | $0.95 \pm 0.002$ / $0.82 \pm 0.014$<br>15.6 m |
| 32 | 8 | $0.95 \pm 0.003$ / $0.70 \pm 0.012$<br>12.7 m | $0.95 \pm 0.002$ / $0.72 \pm 0.015$<br>13.1 m | $0.95 \pm 0.002$ / $0.76 \pm 0.015$<br>14.1 m | $0.95 \pm 0.002$ / $0.83 \pm 0.012$<br>15.7 m |
| 32 | 16 | $0.95 \pm 0.002$ / $0.71 \pm 0.012$<br>13.0 m | $0.95 \pm 0.002$ / $0.74 \pm 0.013$<br>13.4 m | $0.95 \pm 0.002$ / $0.79 \pm 0.011$<br>14.3 m | $0.95 \pm 0.002$ / $0.85 \pm 0.009$<br>15.6 m |
| 32 | 32 | $0.94 \pm 0.002$ / $0.70 \pm 0.012$<br>13.3 m | $0.95 \pm 0.002$ / $0.72 \pm 0.014$<br>13.7 m | $0.95 \pm 0.002$ / $0.79 \pm 0.014$<br>14.2 m | $0.95 \pm 0.002$ / $0.85 \pm 0.010$<br>15.6 m |
| | | **32 Taxa** | | | |
| 8 | 0 | $0.95 \pm 0.002$ / $0.60 \pm 0.015$<br>1.2 m | $0.95 \pm 0.002$ / $0.65 \pm 0.014$<br>1.2 m | $0.95 \pm 0.002$ / $0.69 \pm 0.016$<br>1.4 m | $0.95 \pm 0.002$ / $0.79 \pm 0.011$<br>1.6 m |
| 8 | 8 | $0.95 \pm 0.002$ / $0.63 \pm 0.015$<br>1.2 m | $0.95 \pm 0.002$ / $0.64 \pm 0.015$<br>1.3 m | $0.95 \pm 0.002$ / $0.74 \pm 0.012$<br>1.4 m | $0.95 \pm 0.002$ / $0.78 \pm 0.012$<br>1.7 m |
| 8 | 16 | $0.95 \pm 0.002$ / $0.64 \pm 0.015$<br>1.1 m | $0.95 \pm 0.002$ / $0.66 \pm 0.015$<br>1.2 m | $0.95 \pm 0.001$ / $0.69 \pm 0.017$<br>1.4 m | $0.95 \pm 0.001$ / $0.76 \pm 0.014$<br>1.8 m |

*continued on next page*

*Table 1 continued from previous page*

| | | Individuals per Taxon | | | |
|---|---|---|---|---|---|
| **Traits** | **Missing** | **2** | **4** | **8** | **16** |
| 8 | 32 | $0.95 \pm 0.002$ / $0.62 \pm 0.016$<br>1.2 m | $0.95 \pm 0.001$ / $0.63 \pm 0.017$<br>1.3 m | $0.95 \pm 0.001$ / $0.71 \pm 0.015$<br>1.4 m | $0.95 \pm 0.001$ / $0.76 \pm 0.014$<br>1.8 m |
| 16 | 0 | $0.95 \pm 0.002$ / $0.66 \pm 0.013$<br>3.9 m | $0.95 \pm 0.002$ / $0.70 \pm 0.013$<br>4.1 m | $0.95 \pm 0.001$ / $0.74 \pm 0.013$<br>4.5 m | $0.95 \pm 0.002$ / $0.80 \pm 0.012$<br>5.2 m |
| 16 | 8 | $0.95 \pm 0.002$ / $0.67 \pm 0.013$<br>4.0 m | $0.95 \pm 0.001$ / $0.69 \pm 0.014$<br>4.1 m | $0.95 \pm 0.002$ / $0.75 \pm 0.015$<br>4.5 m | $0.95 \pm 0.001$ / $0.81 \pm 0.009$<br>5.3 m |
| 16 | 16 | $0.95 \pm 0.002$ / $0.64 \pm 0.014$<br>4.0 m | $0.95 \pm 0.002$ / $0.70 \pm 0.012$<br>4.1 m | $0.95 \pm 0.001$ / $0.73 \pm 0.013$<br>4.6 m | $0.95 \pm 0.001$ / $0.80 \pm 0.013$<br>5.3 m |
| 16 | 32 | $0.95 \pm 0.002$ / $0.65 \pm 0.016$<br>4.1 m | $0.95 \pm 0.001$ / $0.67 \pm 0.016$<br>4.3 m | $0.95 \pm 0.001$ / $0.72 \pm 0.014$<br>4.6 m | $0.95 \pm 0.002$ / $0.79 \pm 0.011$<br>5.4 m |
| 32 | 0 | $0.95 \pm 0.002$ / $0.67 \pm 0.014$<br>15.1 m | $0.95 \pm 0.002$ / $0.73 \pm 0.012$<br>15.5 m | $0.95 \pm 0.001$ / $0.77 \pm 0.013$<br>16.0 m | $0.95 \pm 0.001$ / $0.82 \pm 0.009$<br>17.3 m |
| 32 | 8 | $0.94 \pm 0.002$ / $0.68 \pm 0.013$<br>15.0 m | $0.95 \pm 0.001$ / $0.69 \pm 0.017$<br>15.3 m | $0.95 \pm 0.001$ / $0.76 \pm 0.013$<br>16.0 m | $0.95 \pm 0.001$ / $0.79 \pm 0.016$<br>17.3 m |
| 32 | 16 | $0.95 \pm 0.002$ / $0.68 \pm 0.015$<br>15.1 m | $0.95 \pm 0.001$ / $0.70 \pm 0.013$<br>15.4 m | $0.95 \pm 0.001$ / $0.75 \pm 0.013$<br>15.8 m | $0.95 \pm 0.001$ / $0.83 \pm 0.011$<br>17.0 m |
| 32 | 32 | $0.95 \pm 0.002$ / $0.68 \pm 0.013$<br>14.1 m | $0.95 \pm 0.001$ / $0.71 \pm 0.012$<br>14.2 m | $0.95 \pm 0.001$ / $0.74 \pm 0.013$<br>15.1 m | $0.95 \pm 0.001$ / $0.80 \pm 0.012$<br>16.0 m |

*Note.* Each cell reports coverage probability (mean ± SE) for the full model / taxon-means model above the rule, and mean MCMC runtime (minutes per 10 million cycles, full model) below the rule.

| $C_I$ | | | | | | | | |
|---|---|---|---|---|---|---|---|---|
| | Decile 3 | Decile 4 | Decile 5 | Decile 6 | Decile 7 | Decile 8 | Decile 9 | Decile 10 |
| Decile 3 | 95.8 | 97.97 | 96.62 | 98.83 | 94.94 | 87.77 | 85.37 | 89.46 |
| Decile 4 | 97.97 | 92.18 | 90.36 | 93.28 | 91.42 | 83.85 | 83.05 | 84.76 |
| Decile 5 | 96.62 | 90.36 | 85.92 | 85.8 | 81.9 | 71.25 | 71.45 | 73.37 |
| Decile 6 | 98.83 | 93.28 | 85.8 | 84.69 | 75.53 | 64.41 | 64.33 | 67.08 |
| Decile 7 | 94.94 | 91.42 | 81.9 | 75.53 | 64.94 | 52.61 | 51.43 | 54.51 |
| Decile 8 | 87.77 | 83.85 | 71.25 | 64.41 | 52.61 | 43.16 | 40.85 | 44.03 |
| Decile 9 | 85.37 | 83.05 | 71.45 | 64.33 | 51.43 | 40.85 | 40.59 | 42.9 |
| Decile 10 | 89.46 | 84.76 | 73.37 | 67.08 | 54.51 | 44.03 | 42.9 | 47.61 |

| $I^2$ | | | | | | | | |
|---|---|---|---|---|---|---|---|---|
| | Decile 3 | Decile 4 | Decile 5 | Decile 6 | Decile 7 | Decile 8 | Decile 9 | Decile 10 |
| Decile 3 | 92.3 | 93.59 | 95.58 | 90.25 | 93.12 | 93.11 | 92.84 | 93.49 |
| Decile 4 | 93.59 | 94.37 | 91.59 | 88.54 | 84.44 | 83.3 | 83.48 | 87.84 |
| Decile 5 | 95.58 | 91.59 | 89.03 | 78.66 | 75.79 | 70.55 | 70.05 | 72.05 |
| Decile 6 | 90.25 | 88.54 | 78.66 | 77.33 | 75.09 | 74.25 | 73.39 | 73.7 |
| Decile 7 | 93.12 | 84.44 | 75.79 | 75.09 | 75.41 | 70.25 | 70.65 | 73.52 |
| Decile 8 | 93.11 | 83.3 | 70.55 | 74.25 | 70.25 | 70.19 | 68.17 | 71.28 |
| Decile 9 | 92.84 | 83.48 | 70.05 | 73.39 | 70.65 | 68.17 | 68.82 | 70.85 |
| Decile 10 | 93.49 | 87.84 | 72.05 | 73.7 | 73.52 | 71.28 | 70.85 | 73.58 |

**Table 2:** Overlap percentage for each element of the evolutionary variance-covariance matrices between the posterior distributions inferred with the full model and the empirical species means model. Higher overlap indicates the posterior probability distribution for that element of the evolutionary VCV matrix is more similar between the two models.

| Analysis | Modern human $\sigma^2_\mu$ | Neandertal $\sigma^2_\mu$ | *Pan paniscus* $\sigma^2_\mu$ | *Pan troglodytes* $\sigma^2_\mu$ | *G. beringei* $\sigma^2_\mu$ | *G. gorilla* $\sigma^2_\mu$ | *Po. abelii* $\sigma^2_\mu$ | *Po. pygmaeus* $\sigma^2_\mu$ |
|---|---|---|---|---|---|---|---|---|
| $C_1$ | 10.19 | 1.56 | 3.84 | 2.52 | 2.53 | 4.14 | 4.37 | 4.22 |
| $I^2$ | 5.48 | 3.74 | 58.37 | 6.16 | - | - | - | - |

**Table 3:** Total variance in the posterior distribution for the inferred intraspecific means ($\sigma^2_\mu$). Calculated as the sum across all deciles of the variance in the posterior distribution for that decile.

| Analysis | Modern human VCV | Neandertal VCV | *Pan paniscus* VCV | *Pan troglodytes* VCV | *G. beringei* VCV | *G. gorilla* VCV | *Po. abelii* VCV | *Po. pygmaeus* VCV |
|---|---|---|---|---|---|---|---|---|
| $C_1$ | 3,739.98 | 510.46 | 2,152.99 | 805.53 | 291.81 | 139.11 | 142.1 | 322.09 |
| $I^2$ | 7,400.58 | 276.37 | 66.28 | 577.19 | - | - | - | - |

**Table 4:** Symmetrized Kullback-Leibler divergences measuring the divergence in the inferred evolutionary VCV matrix with each intraspecific VCV matrix.

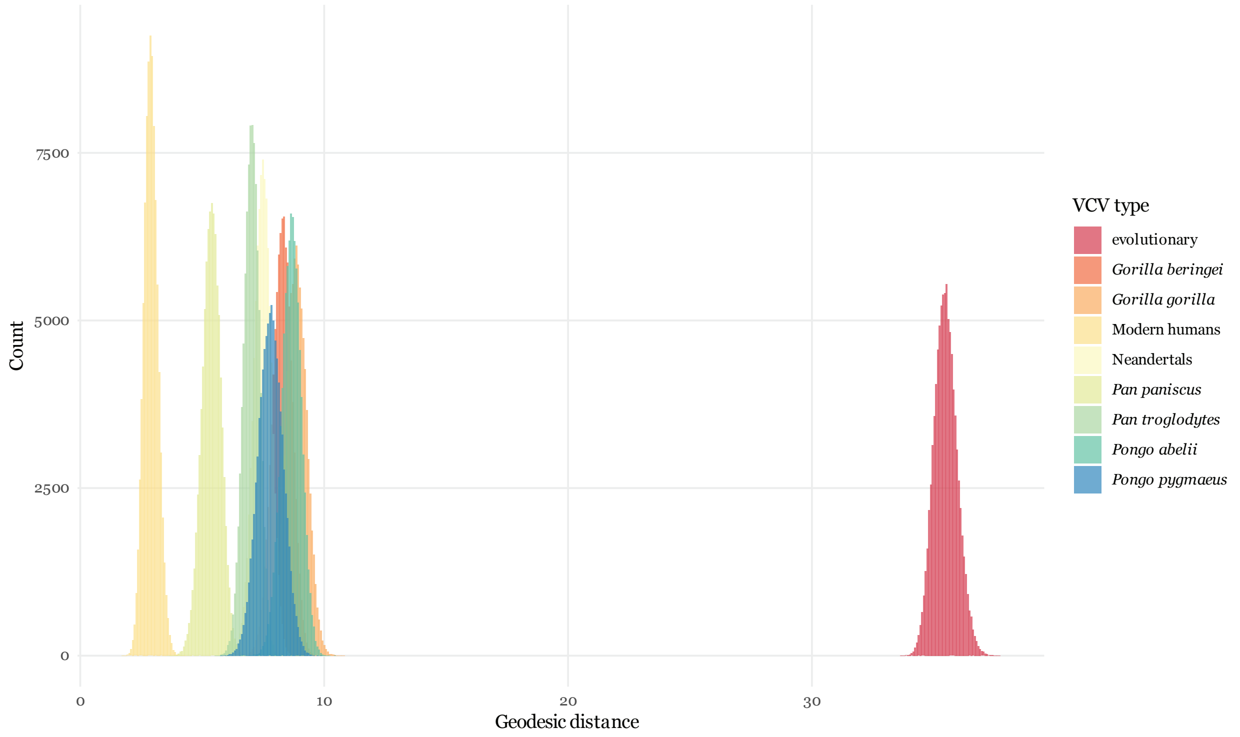


**Figure 1: Geodesic distances between MLE estimates and the posterior distribution for each VCV matrix.** We calculated the geodesic distances between each VCV matrix in our $C_1$ posterior distribution to the MLE singular intraspecific VCV and evolutionary VCV under the Felsenstein (Felsenstein, 2008) model as implemented in *rphylopars* (Goolsby et al., 2017). A distance of zero indicates that the two matrices are identical. Histograms were built with 500 bins.

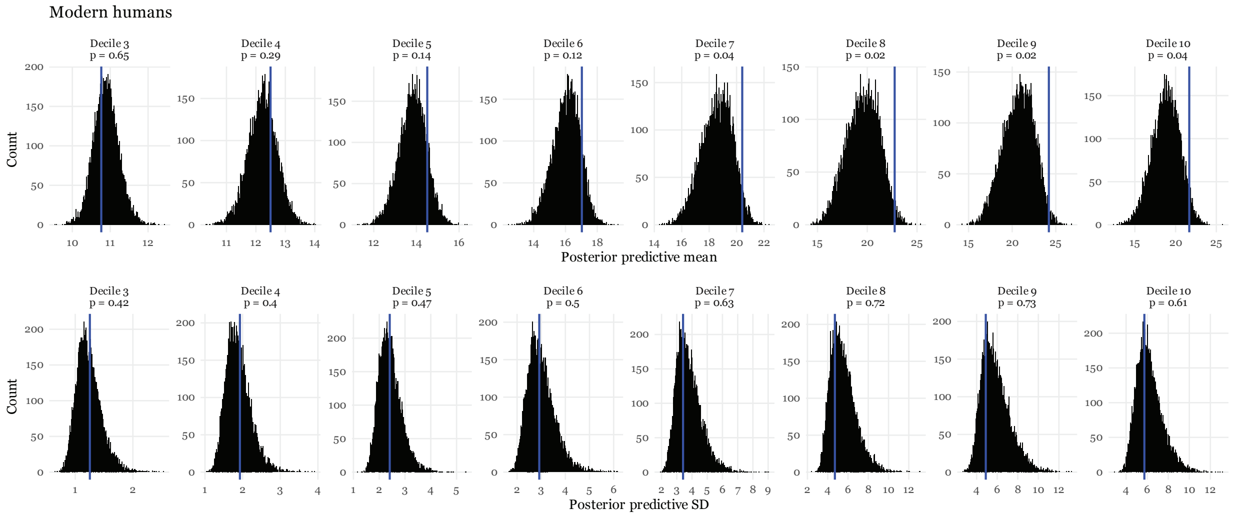


**Figure 2: Posterior predictive mean and standard deviations for the modern human $C_1$ data.** The horizontal axis indicates the posterior predictive summary statistic, either mean (top row) or standard deviation (bottom row). Above each facet is the Bayesian *p*-value, indicating the proportion of posterior predictive test statistics that exceed the empirically observed test statistic; *p*-values close to 0 or 1 indicate poor fit for that decile. The vertical axis describes the frequency of that summary statistic among posterior predictive summary statistics. The blue vertical line indicates the empirical mean or standard deviation for each decile. Histograms were built with 500 bins.

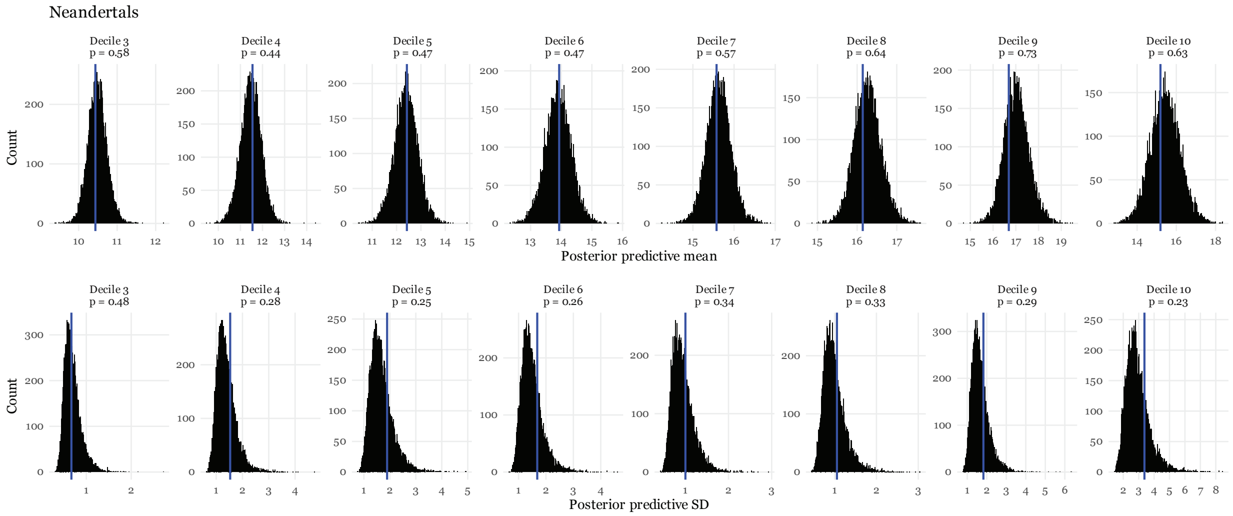


**Figure 3: Posterior predictive mean and standard deviations for the Neandertal $C_1$ data.** The horizontal axis indicates the posterior predictive summary statistic, either mean (top row) or standard deviation (bottom row). Above each facet is the Bayesian *p*-value, indicating the proportion of posterior predictive test statistics that exceed the empirically observed test statistic; *p*-values close to 0 or 1 indicate poor fit for that decile. The vertical axis describes the frequency of that summary statistic among posterior predictive summary statistics. The blue vertical line indicates the empirical mean or standard deviation for each decile. Histograms were built with 500 bins.

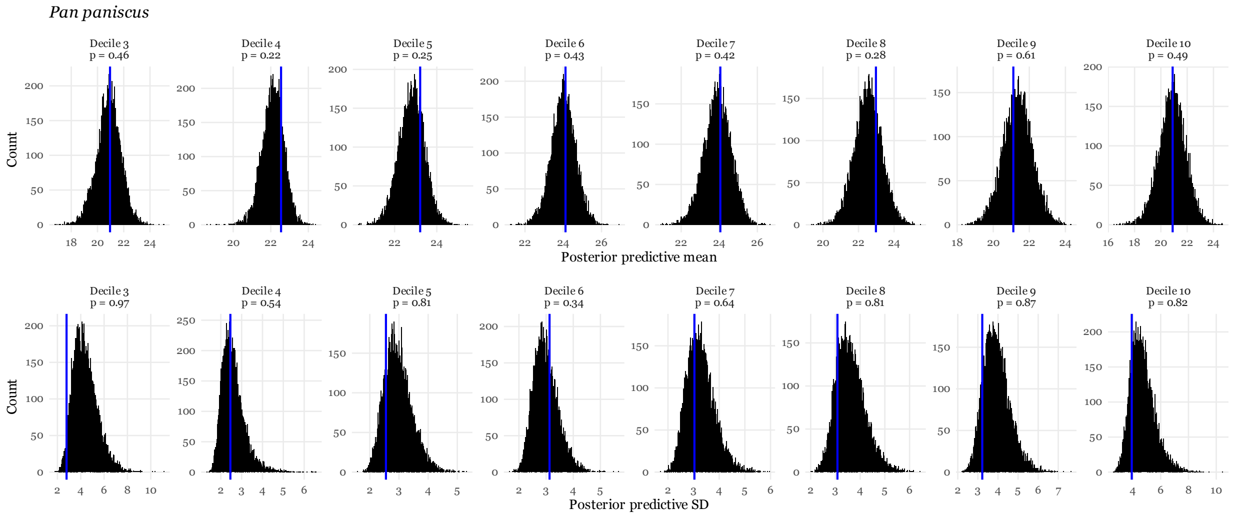


**Figure 4: Posterior predictive mean and standard deviations for the *Pan paniscus* $C_1$ data.** The horizontal axis indicates the posterior predictive summary statistic, either mean (top row) or standard deviation (bottom row). Above each facet is the Bayesian *p*-value, indicating the proportion of posterior predictive test statistics that exceed the empirically observed test statistic; *p*-values close to 0 or 1 indicate poor fit for that decile. The vertical axis describes the frequency of that summary statistic among posterior predictive summary statistics. The blue vertical line indicates the empirical mean or standard deviation for each decile. Histograms were built with 500 bins.

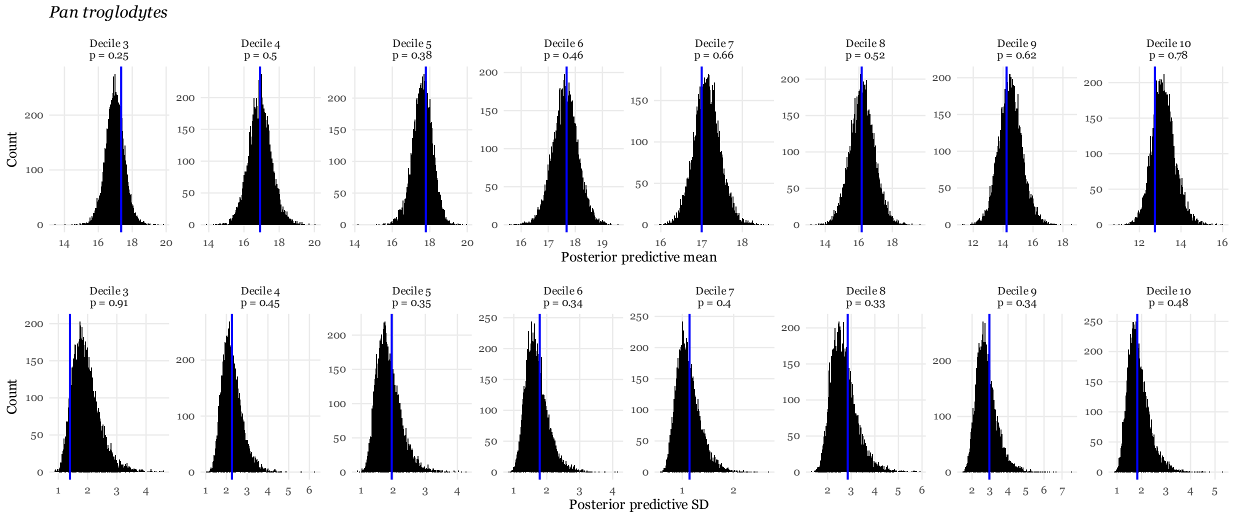


**Figure 5: Posterior predictive mean and standard deviations for the *Pan troglodytes* $C_1$ data.** The horizontal axis indicates the posterior predictive summary statistic, either mean (top row) or standard deviation (bottom row). Above each facet is the Bayesian *p*-value, indicating the proportion of posterior predictive test statistics that exceed the empirically observed test statistic; *p*-values close to 0 or 1 indicate poor fit for that decile. The vertical axis describes the frequency of that summary statistic among posterior predictive summary statistics. The blue vertical line indicates the empirical mean or standard deviation for each decile. Histograms were built with 500 bins.

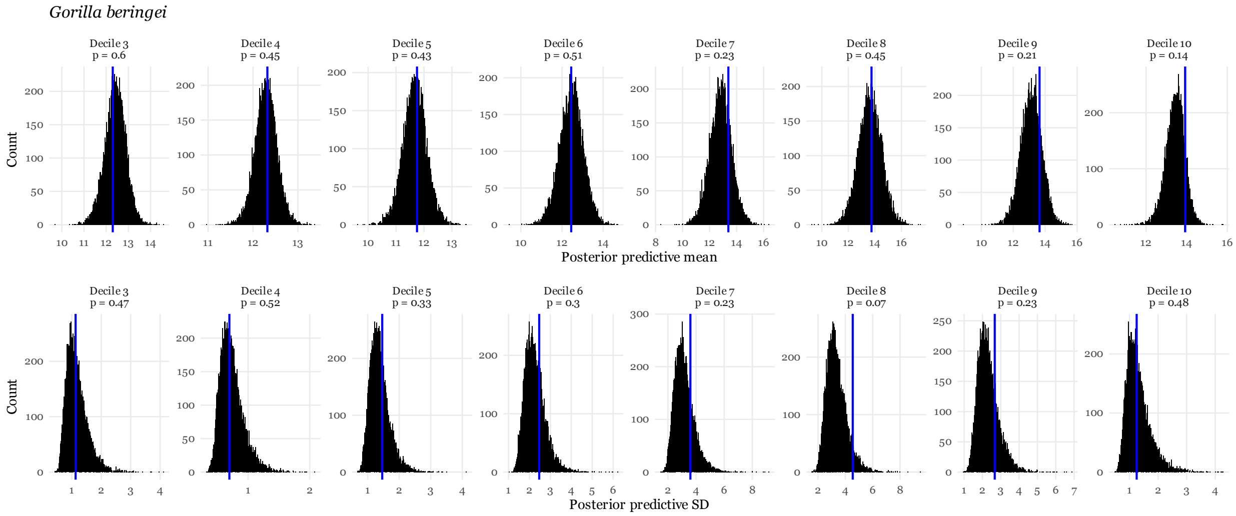


**Figure 6: Posterior predictive mean and standard deviations for the *Gorilla beringei* $C_1$ data.** The horizontal axis indicates the posterior predictive summary statistic, either mean (top row) or standard deviation (bottom row). Above each facet is the Bayesian *p*-value, indicating the proportion of posterior predictive test statistics that exceed the empirically observed test statistic; *p*-values close to 0 or 1 indicate poor fit for that decile. The vertical axis describes the frequency of that summary statistic among posterior predictive summary statistics. The blue vertical line indicates the empirical mean or standard deviation for each decile. Histograms were built with 500 bins.

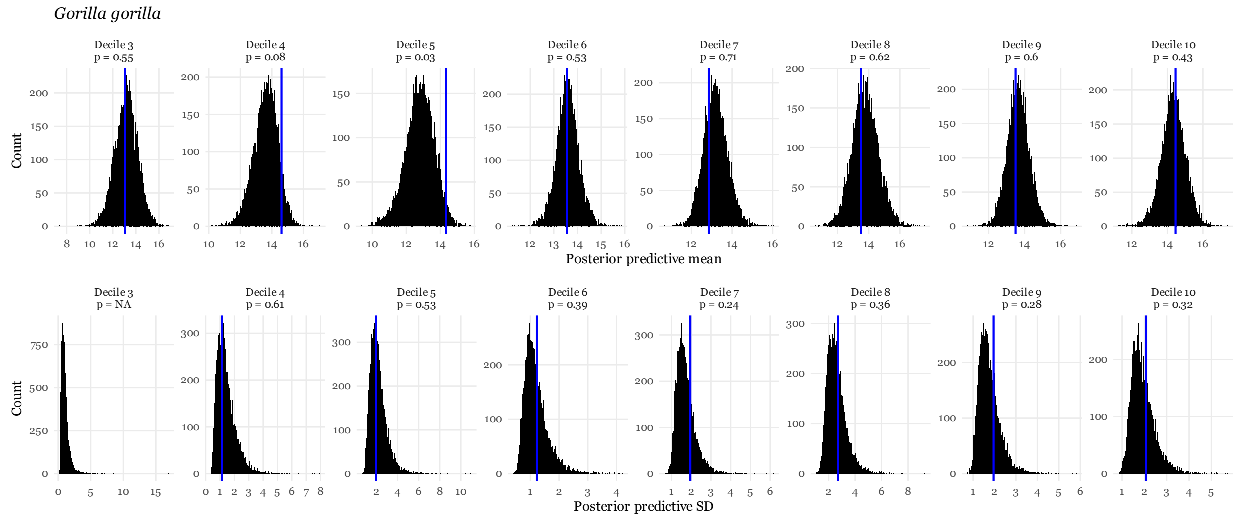


**Figure 7: Posterior predictive mean and standard deviations for the *Gorilla gorilla* $C_1$ data.** The horizontal axis indicates the posterior predictive summary statistic, either mean (top row) or standard deviation (bottom row). Above each facet is the Bayesian *p*-value, indicating the proportion of posterior predictive test statistics that exceed the empirically observed test statistic; *p*-values close to 0 or 1 indicate poor fit for that decile. The vertical axis describes the frequency of that summary statistic among posterior predictive summary statistics. The blue vertical line indicates the empirical mean or standard deviation for each decile. As we only have one sample for the *G. gorilla* decile 3 (all other individuals had that decile missing), we were unable to calculate the SD for that decile, hence the absent blue line. Histograms were built with 500 bins.

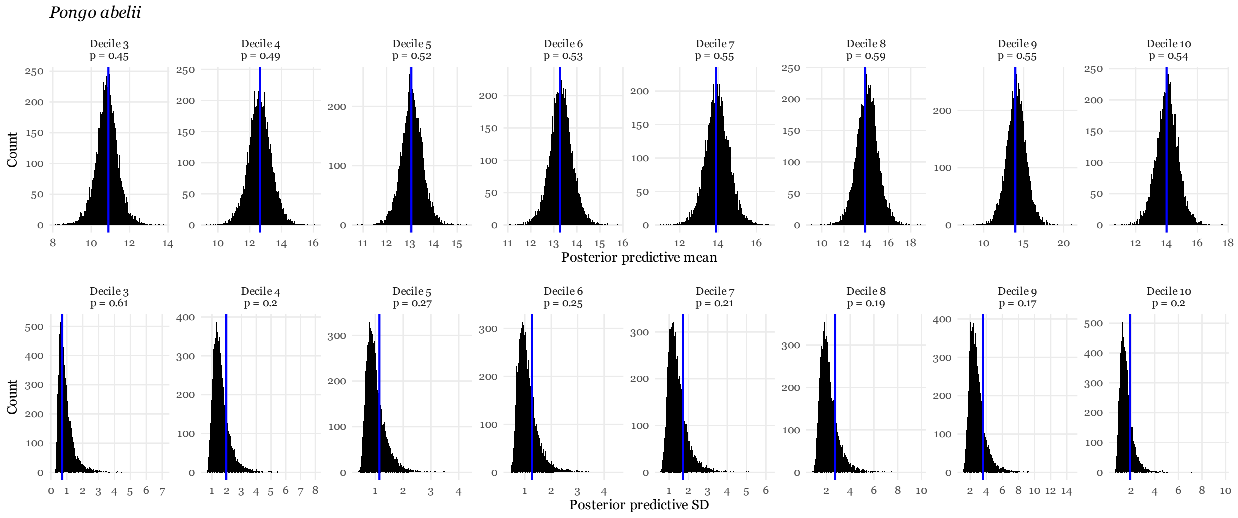


**Figure 8: Posterior predictive mean and standard deviations for the *Pongo abelii* $C_1$ data.** The horizontal axis indicates the posterior predictive summary statistic, either mean (top row) or standard deviation (bottom row). Above each facet is the Bayesian *p*-value, indicating the proportion of posterior predictive test statistics that exceed the empirically observed test statistic; *p*-values close to 0 or 1 indicate poor fit for that decile. The vertical axis describes the frequency of that summary statistic among posterior predictive summary statistics. The blue vertical line indicates the empirical mean or standard deviation for each decile. Histograms were built with 500 bins.

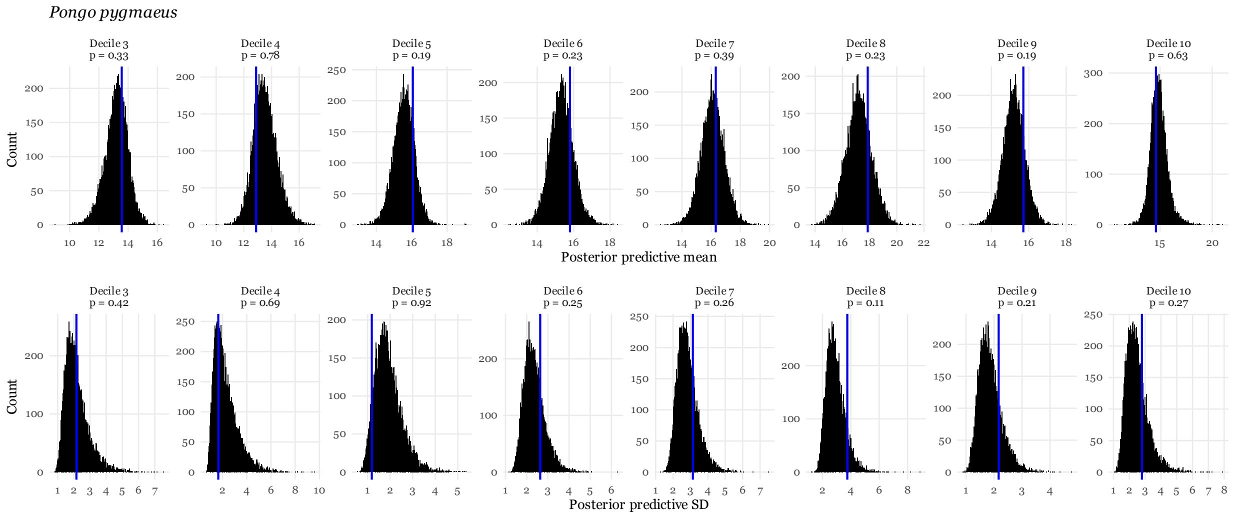


**Figure 9: Posterior predictive mean and standard deviations for the *Pongo pygmaeus* $C_1$ data.** The horizontal axis indicates the posterior predictive summary statistic, either mean (top row) or standard deviation (bottom row). Above each facet is the Bayesian *p*-value, indicating the proportion of posterior predictive test statistics that exceed the empirically observed test statistic; *p*-values close to 0 or 1 indicate poor fit for that decile. The vertical axis describes the frequency of that summary statistic among posterior predictive summary statistics. The blue vertical line indicates the empirical mean or standard deviation for each decile. Histograms were built with 500 bins.

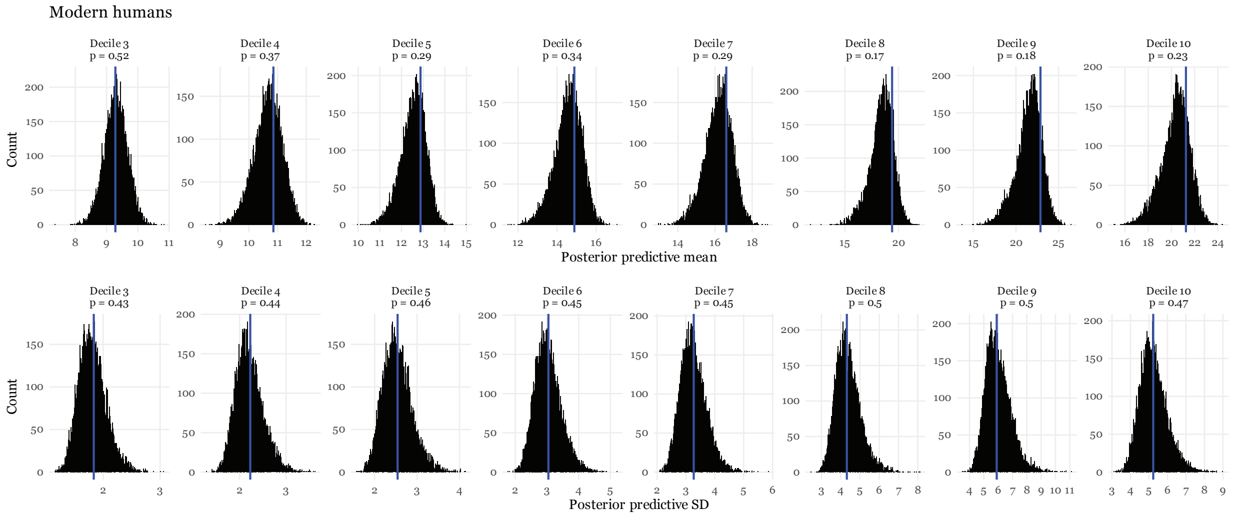


**Figure 10: Posterior predictive mean and standard deviations for the modern human $I^2$ data.** The horizontal axis indicates the posterior predictive summary statistic, either mean (top row) or standard deviation (bottom row). Above each facet is the Bayesian *p*-value, indicating the proportion of posterior predictive test statistics that exceed the empirically observed test statistic; *p*-values close to 0 or 1 indicate poor fit for that decile. The vertical axis describes the frequency of that summary statistic among posterior predictive summary statistics. The blue vertical line indicates the empirical mean or standard deviation for each decile. Histograms were built with 500 bins.

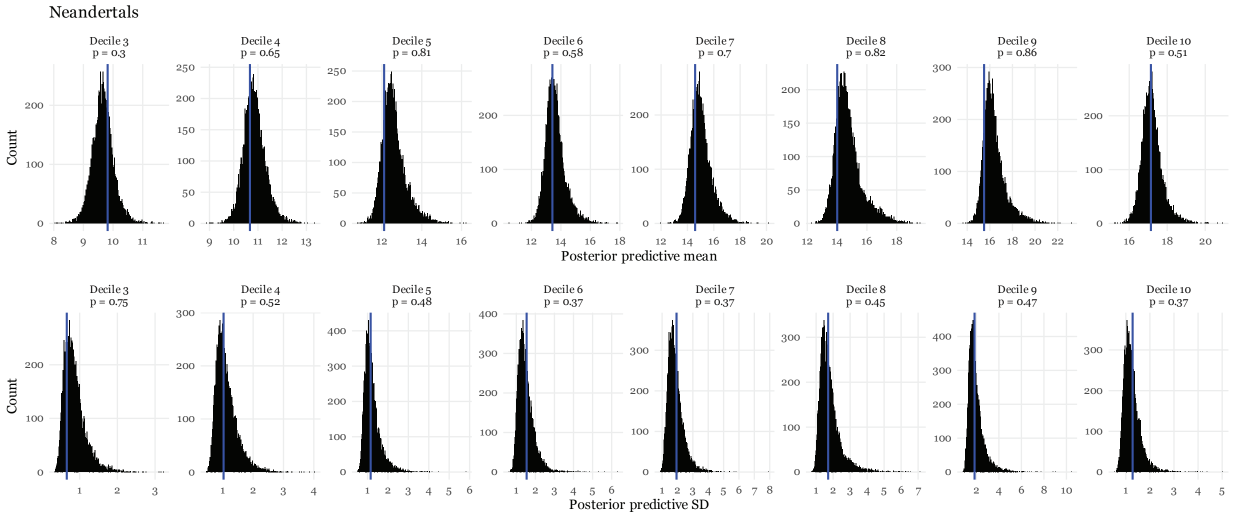


**Figure 11: Posterior predictive mean and standard deviations for the Neandertal I$^2$ data.** The horizontal axis indicates the posterior predictive summary statistic, either mean (top row) or standard deviation (bottom row). Above each facet is the Bayesian *p*-value, indicating the proportion of posterior predictive test statistics that exceed the empirically observed test statistic; *p*-values close to 0 or 1 indicate poor fit for that decile. The vertical axis describes the frequency of that summary statistic among posterior predictive summary statistics. The blue vertical line indicates the empirical mean or standard deviation for each decile. Histograms were built with 500 bins.

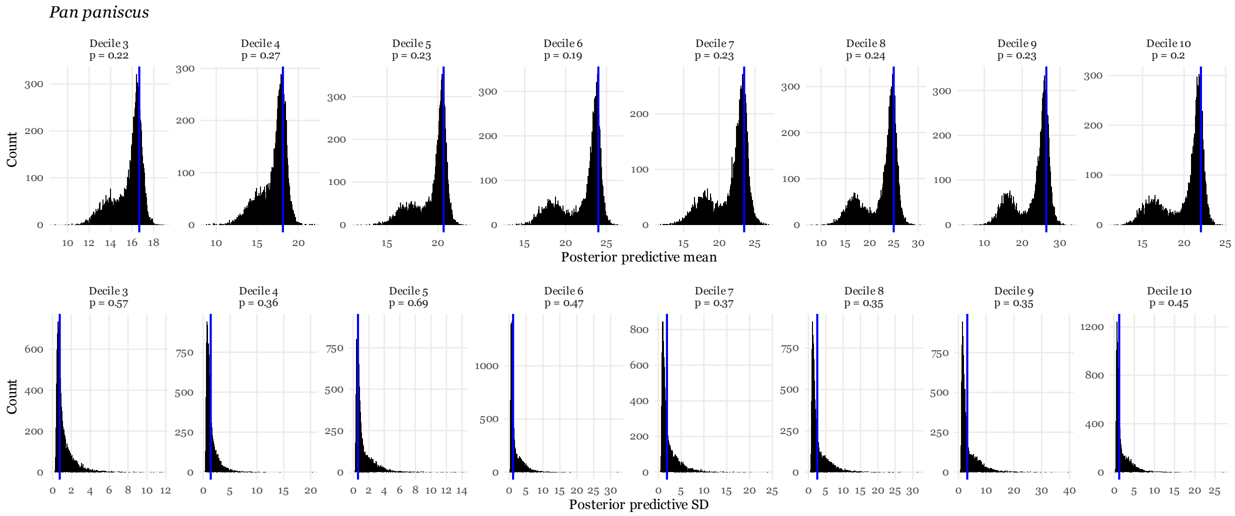


**Figure 12: Posterior predictive mean and standard deviations for the *Pan paniscus* $I^2$ data.** The horizontal axis indicates the posterior predictive summary statistic, either mean (top row) or standard deviation (bottom row). Above each facet is the Bayesian *p*-value, indicating the proportion of posterior predictive test statistics that exceed the empirically observed test statistic; *p*-values close to 0 or 1 indicate poor fit for that decile. The vertical axis describes the frequency of that summary statistic among posterior predictive summary statistics. The blue vertical line indicates the empirical mean or standard deviation for each decile. Histograms were built with 500 bins.

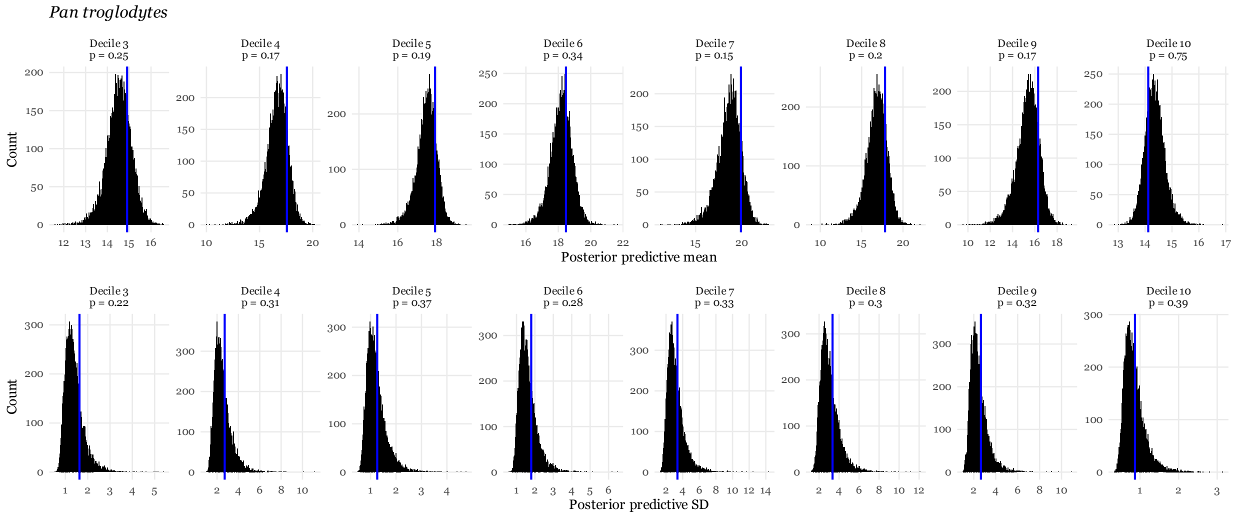


**Figure 13: Posterior predictive mean and standard deviations for the *Pan troglodytes* I$^2$ data.** The horizontal axis indicates the posterior predictive summary statistic, either mean (top row) or standard deviation (bottom row). Above each facet is the Bayesian *p*-value, indicating the proportion of posterior predictive test statistics that exceed the empirically observed test statistic; *p*-values close to 0 or 1 indicate poor fit for that decile. The vertical axis describes the frequency of that summary statistic among posterior predictive summary statistics. The blue vertical line indicates the empirical mean or standard deviation for each decile. Histograms were built with 500 bins.

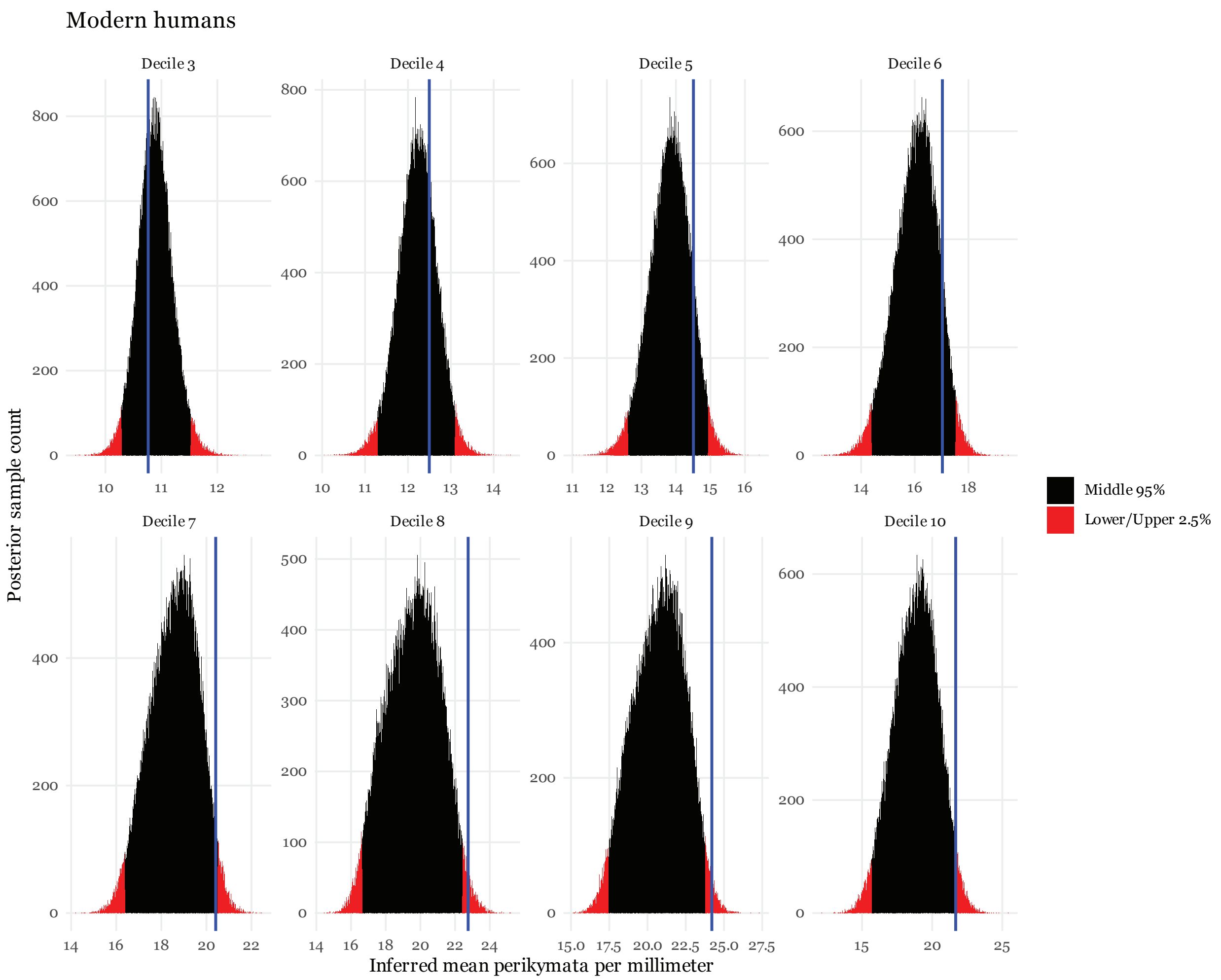


**Figure 14: Posterior distribution for the modern human intraspecific $C_1$ mean**. The horizontal axis indicates the inferred mean perikymata per millimeter for each decile and is truncated between 0 and 30. The vertical axis describes the frequency of that value in the posterior samples. The blue vertical line indicates the empirical mean perikymata per millimeter for each decile. Red bins indicate the lower and upper 2.5% quantiles and the remaining black bins fall within the 95% central credible region. The width of this distribution indicates the statistical uncertainty in the value of this parameter; wider distributions are more uncertain. Histograms were built with 500 bins.

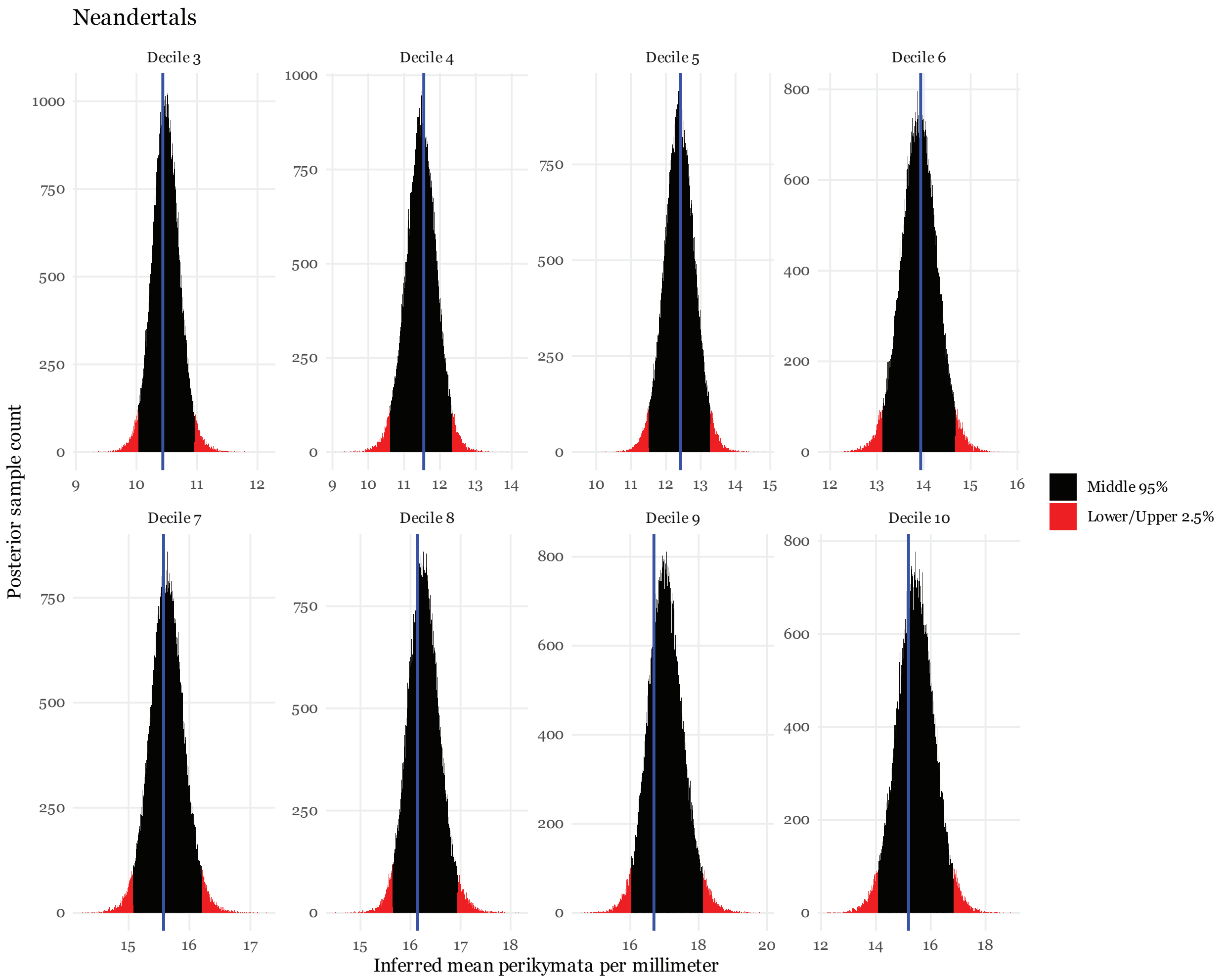


**Figure 15: Posterior distribution for the Neandertal intraspecific $C_1$ mean.** The horizontal axis indicates the inferred mean perikymata per millimeter for each decile and is truncated between 0 and 30. The vertical axis describes the frequency of that value in the posterior samples. The blue vertical line indicates the empirical mean perikymata per millimeter for each decile. Red bins indicate the lower and upper 2.5% quantiles and the remaining black bins fall within the 95% central credible region. The width of this distribution indicates the statistical uncertainty in the value of this parameter; wider distributions are more uncertain. Histograms were built with 500 bins.

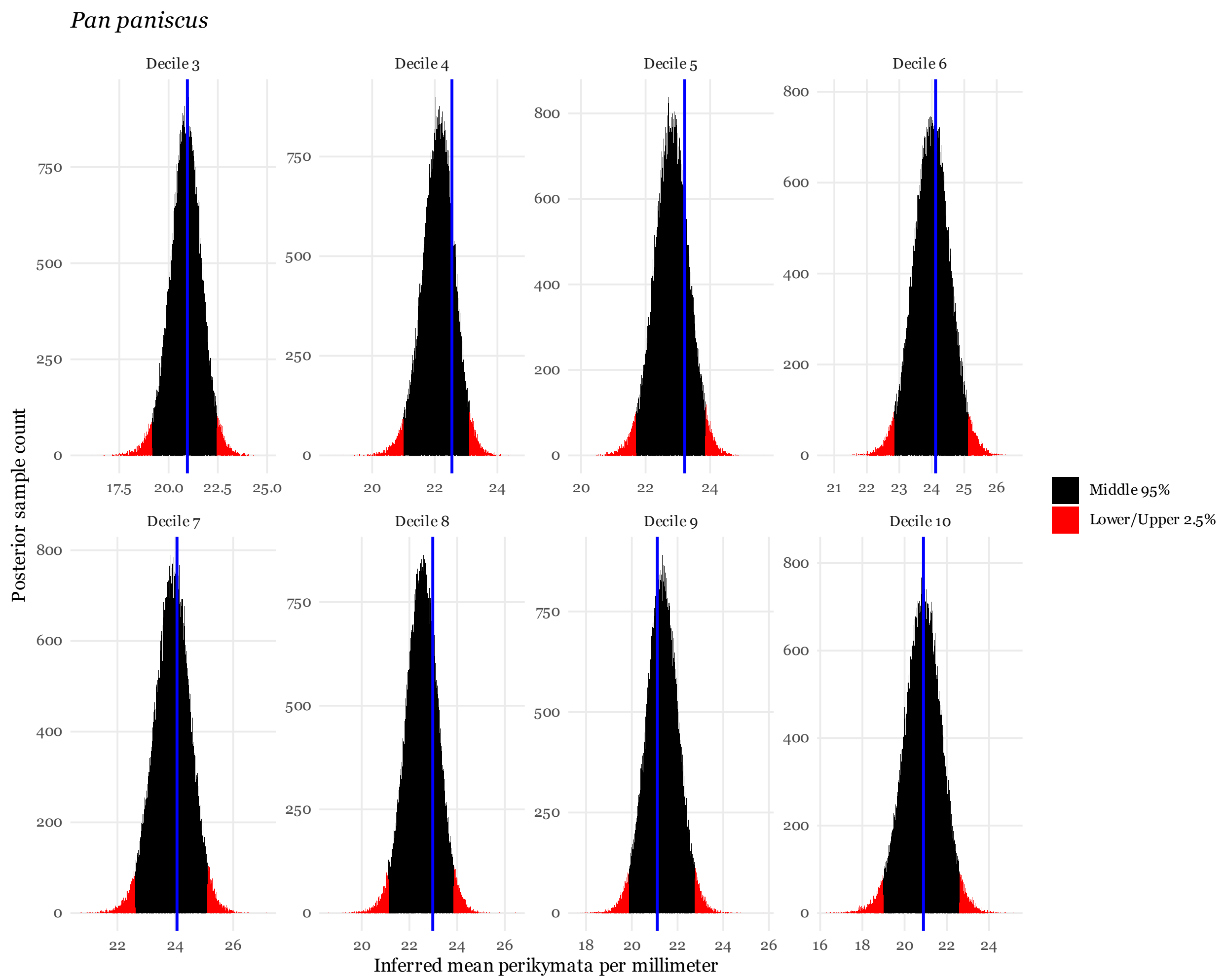


**Figure 16: Posterior distribution for the *Pan paniscus* intraspecific $C_1$ mean.** The horizontal axis indicates the inferred mean perikymata per millimeter for each decile and is truncated between 0 and 30. The vertical axis describes the frequency of that value in the posterior samples. The blue vertical line indicates the empirical mean perikymata per millimeter for each decile. Red bins indicate the lower and upper 2.5% quantiles and the remaining black bins fall within the 95% central credible region. The width of this distribution indicates the statistical uncertainty in the value of this parameter; wider distributions are more uncertain. Histograms were built with 500 bins.

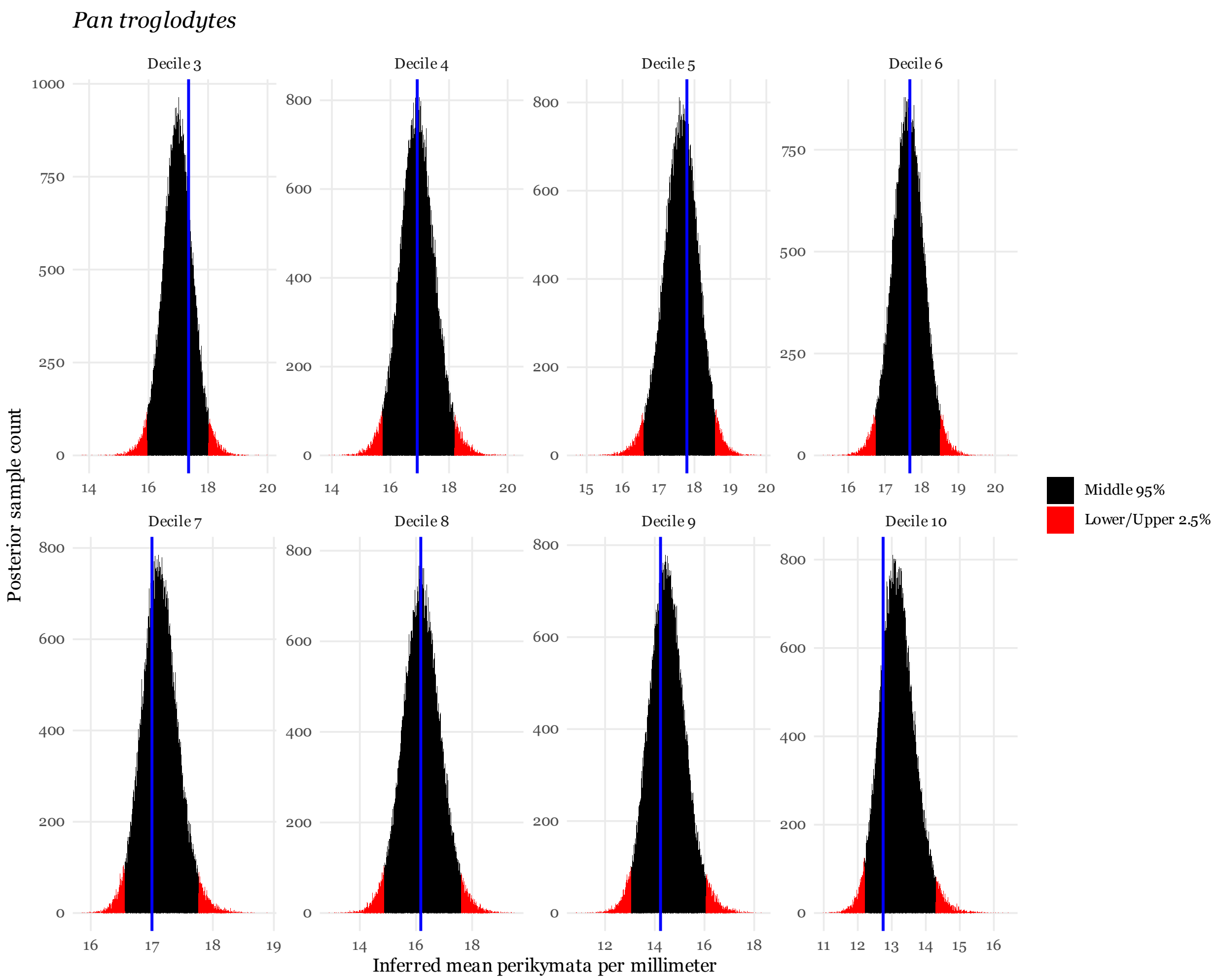


**Figure 17: Posterior distribution for the *Pan troglodytes* intraspecific $C_1$ mean.** The horizontal axis indicates the inferred mean perikymata per millimeter for each decile and is truncated between 0 and 30. The vertical axis describes the frequency of that value in the posterior samples. The blue vertical line indicates the empirical mean perikymata per millimeter for each decile. Red bins indicate the lower and upper 2.5% quantiles and the remaining black bins fall within the 95% central credible region. The width of this distribution indicates the statistical uncertainty in the value of this parameter; wider distributions are more uncertain. Histograms were built with 500 bins.

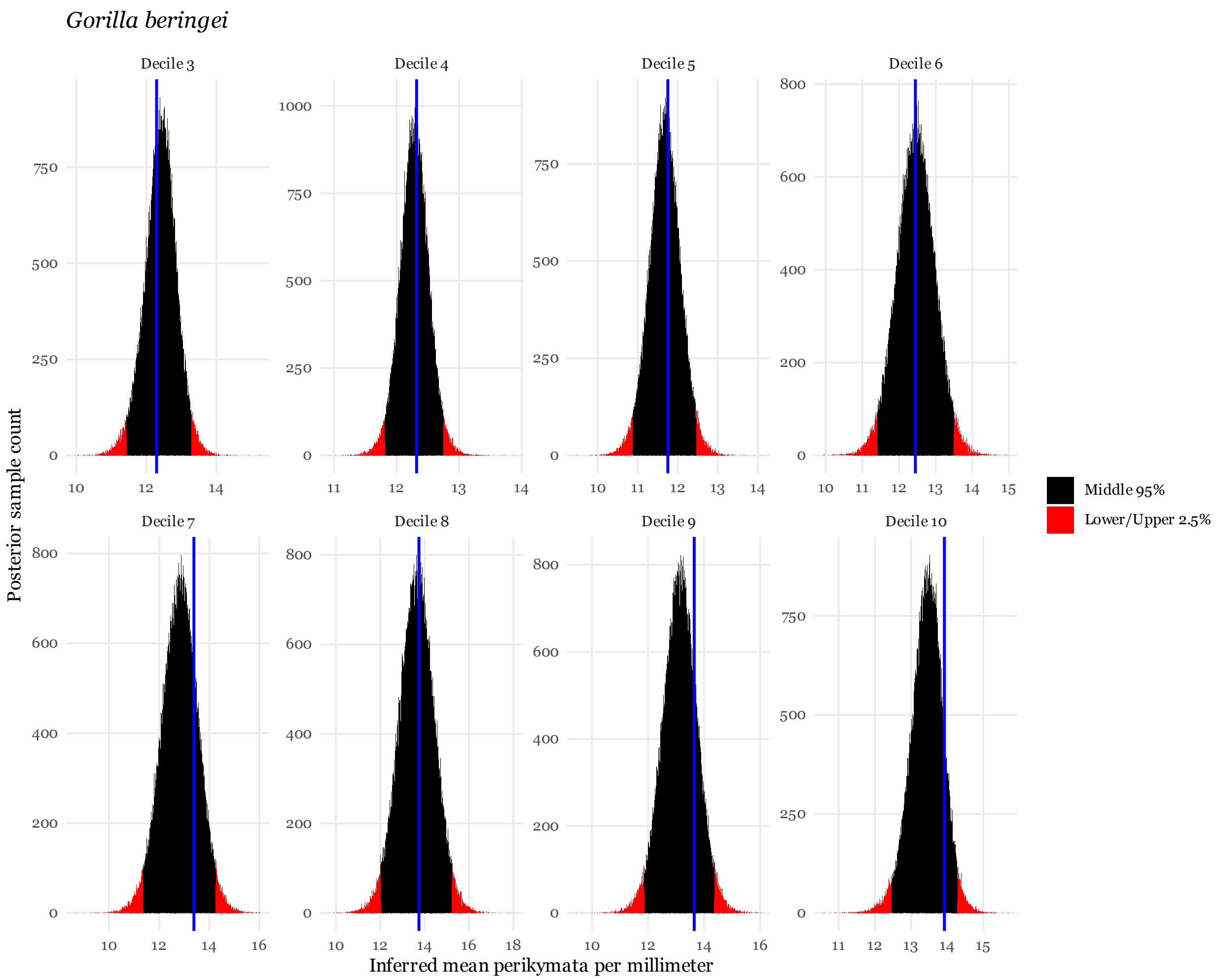


**Figure 18: Posterior distribution for the *Gorilla beringei* intraspecific $C_1$ mean.** The horizontal axis indicates the inferred mean perikymata per millimeter for each decile and is truncated between 0 and 30. The vertical axis describes the frequency of that value in the posterior samples. The blue vertical line indicates the empirical mean perikymata per millimeter for each decile. Red bins indicate the lower and upper 2.5% quantiles and the remaining black bins fall within the 95% central credible region. The width of this distribution indicates the statistical uncertainty in the value of this parameter; wider distributions are more uncertain. Histograms were built with 500 bins.

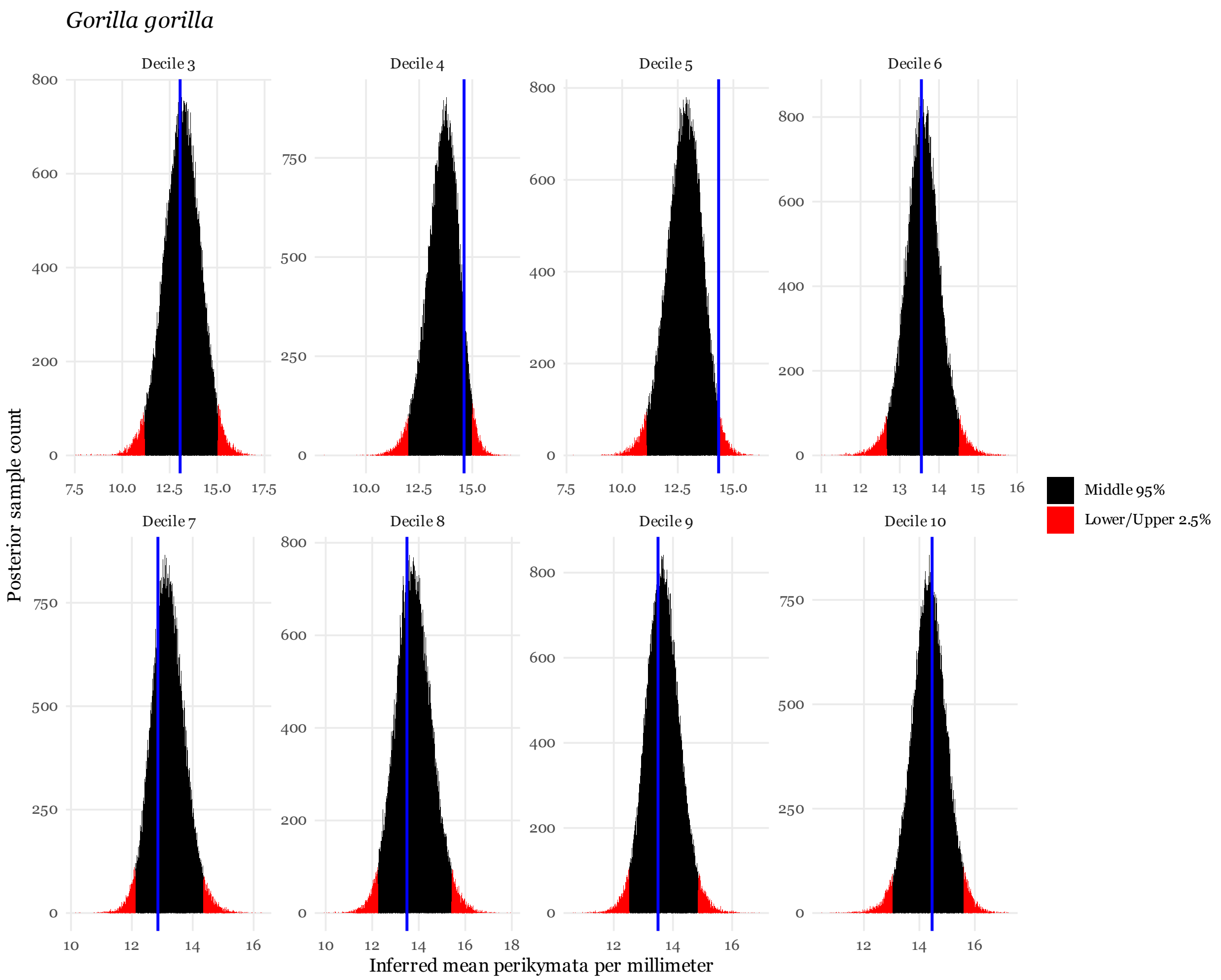


**Figure 19: Posterior distribution for the *Gorilla gorilla* intraspecific $C_1$ mean.** The horizontal axis indicates the inferred mean perikymata per millimeter for each decile and is truncated between 0 and 30. The vertical axis describes the frequency of that value in the posterior samples. The blue vertical line indicates the empirical mean perikymata per millimeter for each decile. Red bins indicate the lower and upper 2.5% quantiles and the remaining black bins fall within the 95% central credible region. The width of this distribution indicates the statistical uncertainty in the value of this parameter; wider distributions are more uncertain. Histograms were built with 500 bins.

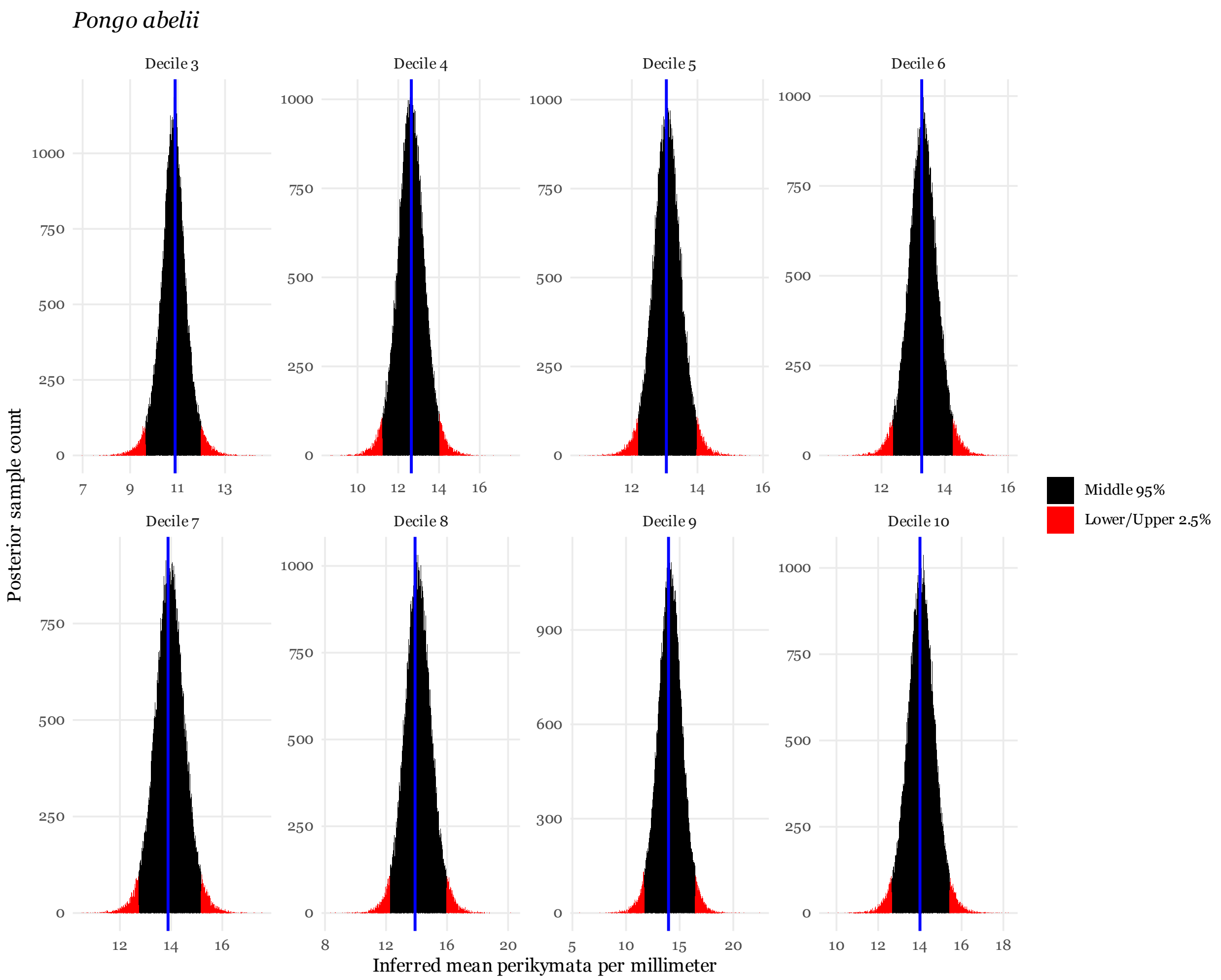


**Figure 20: Posterior distribution for the *Pongo abelii* intraspecific $C_1$ mean.** The horizontal axis indicates the inferred mean perikymata per millimeter for each decile and is truncated between 0 and 30. The vertical axis describes the frequency of that value in the posterior samples. The blue vertical line indicates the empirical mean perikymata per millimeter for each decile. Red bins indicate the lower and upper 2.5% quantiles and the remaining black bins fall within the 95% central credible region. The width of this distribution indicates the statistical uncertainty in the value of this parameter; wider distributions are more uncertain. Histograms were built with 500 bins.

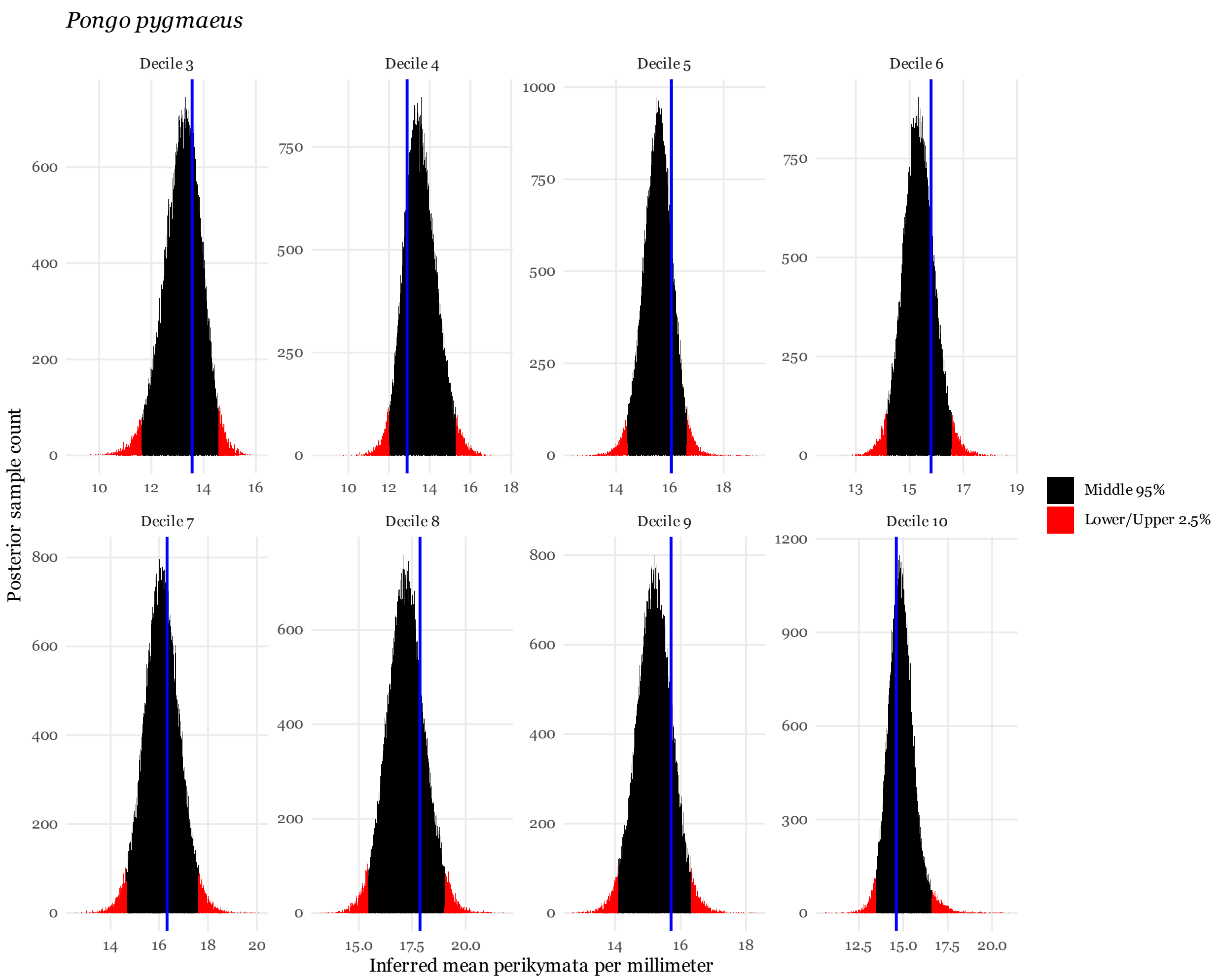


**Figure 21: Posterior distribution for the *Pongo pygmaeus* intraspecific $C_1$ mean.** The horizontal axis indicates the inferred mean perikymata per millimeter for each decile and is truncated between 0 and 30. The vertical axis describes the frequency of that value in the posterior samples. The blue vertical line indicates the empirical mean perikymata per millimeter for each decile. Red bins indicate the lower and upper 2.5% quantiles and the remaining black bins fall within the 95% central credible region. The width of this distribution indicates the statistical uncertainty in the value of this parameter; wider distributions are more uncertain. Histograms were built with 500 bins.

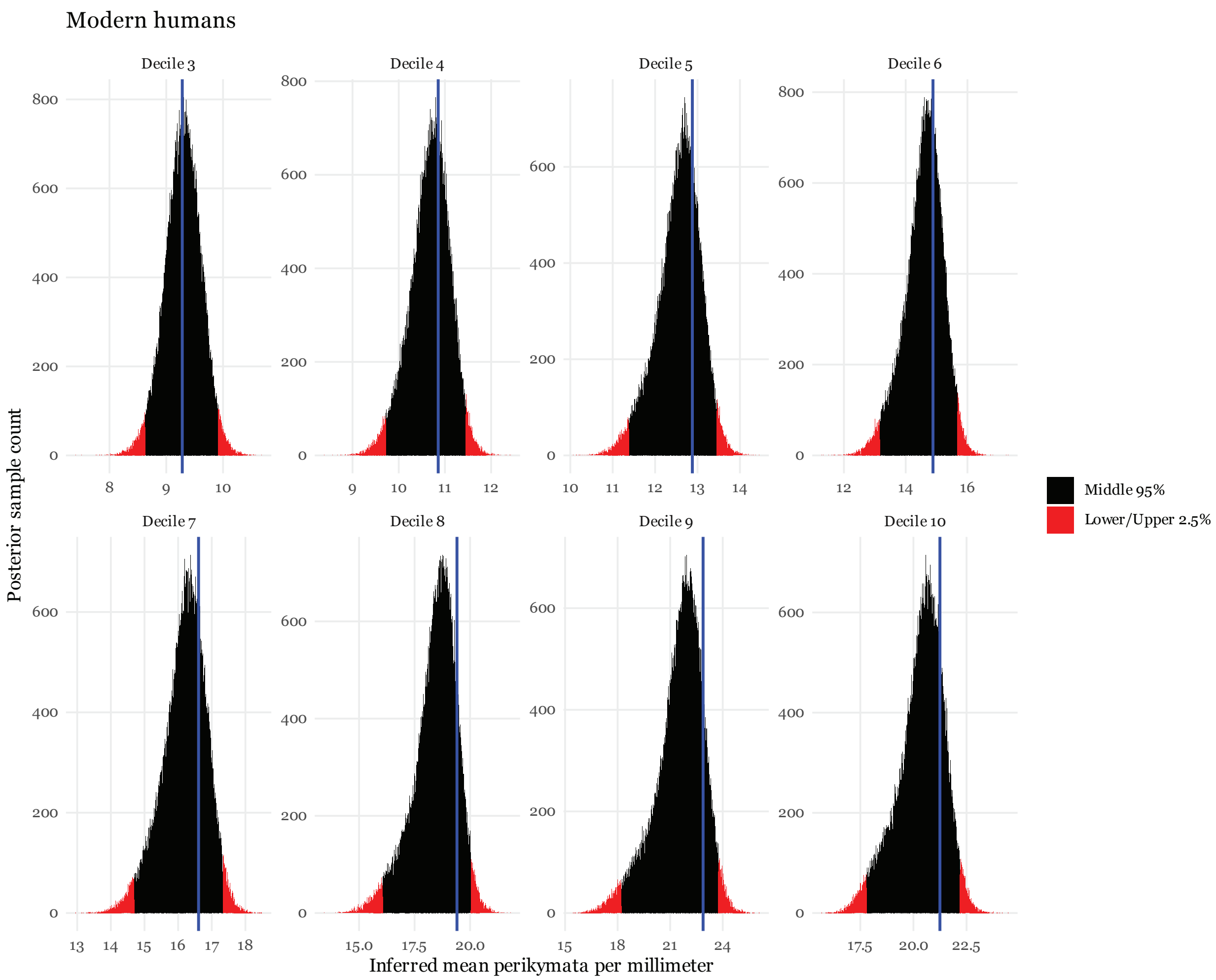


**Figure 22: Posterior distribution for the modern human intraspecific I² mean.** The horizontal axis indicates the inferred mean perikymata per millimeter for each decile and is truncated between 0 and 30. The vertical axis describes the frequency of that value in the posterior samples. The blue vertical line indicates the empirical mean perikymata per millimeter for each decile. Red bins indicate the lower and upper 2.5% quantiles and the remaining black bins fall within the 95% central credible region. The width of this distribution indicates the statistical uncertainty in the value of this parameter; wider distributions are more uncertain. Histograms were built with 500 bins.

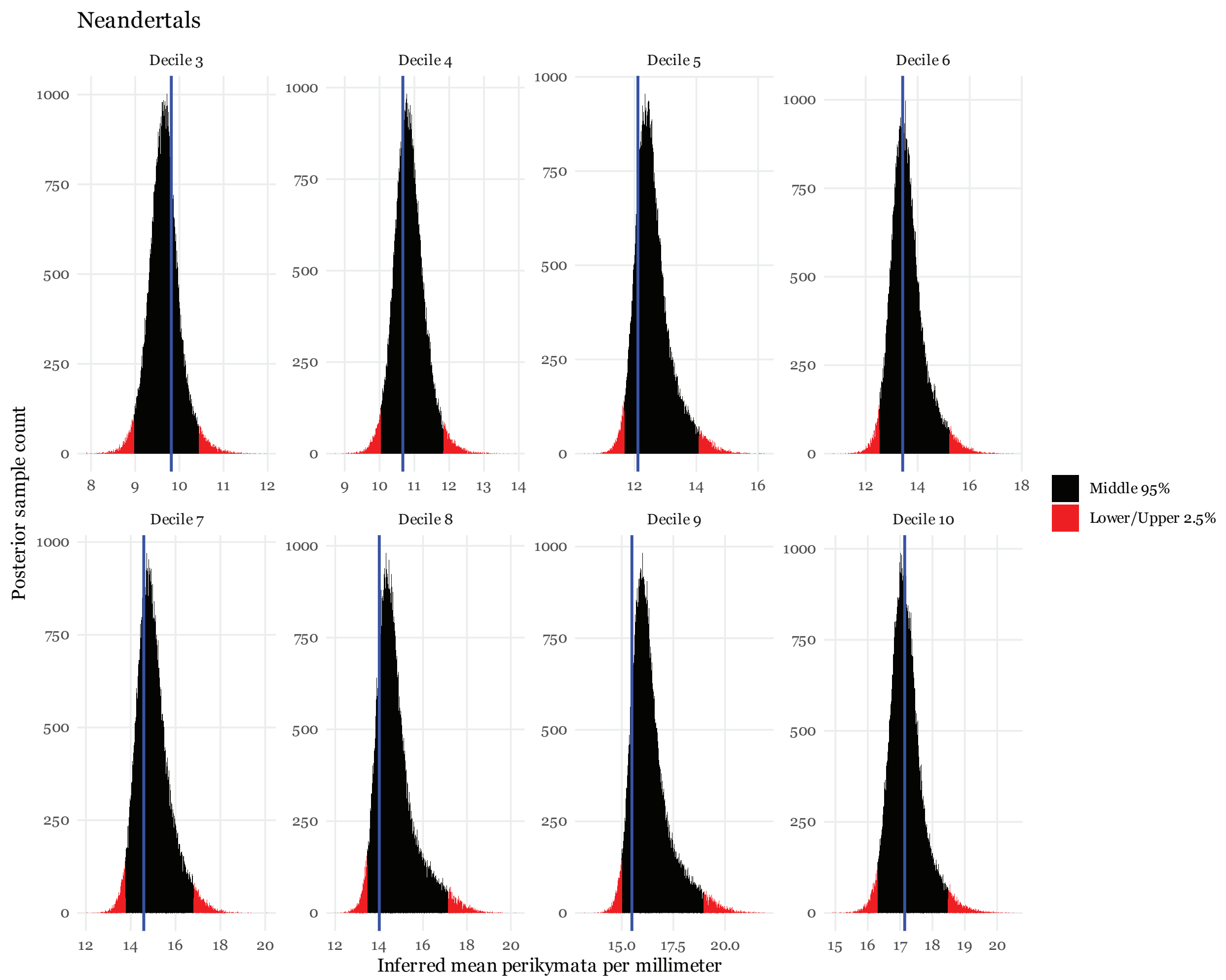


**Figure 23: Posterior distribution for the Neandertal intraspecific $I^2$ mean.** The horizontal axis indicates the inferred mean perikymata per millimeter for each decile and is truncated between 0 and 30. The vertical axis describes the frequency of that value in the posterior samples. The blue vertical line indicates the empirical mean perikymata per millimeter for each decile. Red bins indicate the lower and upper 2.5% quantiles and the remaining black bins fall within the 95% central credible region. The width of this distribution indicates the statistical uncertainty in the value of this parameter; wider distributions are more uncertain. Histograms were built with 500 bins.

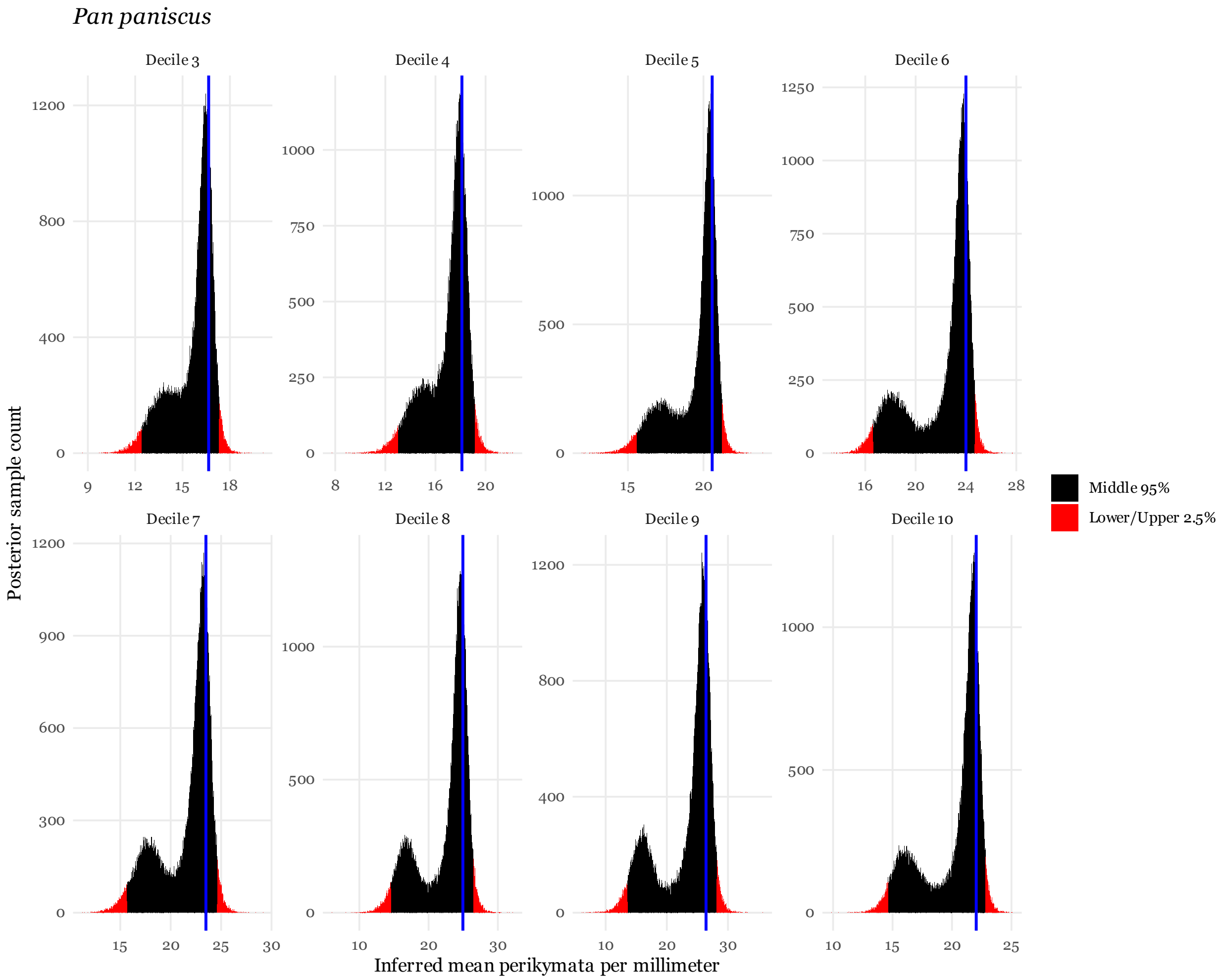


**Figure 24: Posterior distribution for the *Pan paniscus* intraspecific I² mean.** The horizontal axis indicates the inferred mean perikymata per millimeter for each decile. We chose an upper bound of 40 due to the large and relatively diffuse posterior distribution for the decile nine mean. The vertical axis describes the frequency of that value in the posterior samples. The blue vertical line indicates the empirical mean perikymata per millimeter for each decile. Red bins indicate the lower and upper 2.5% quantiles and the remaining black bins fall within the 95% central credible region. The width of this distribution indicates the statistical uncertainty in the value of this parameter; wider distributions are more uncertain. Histograms were built with 500 bins.

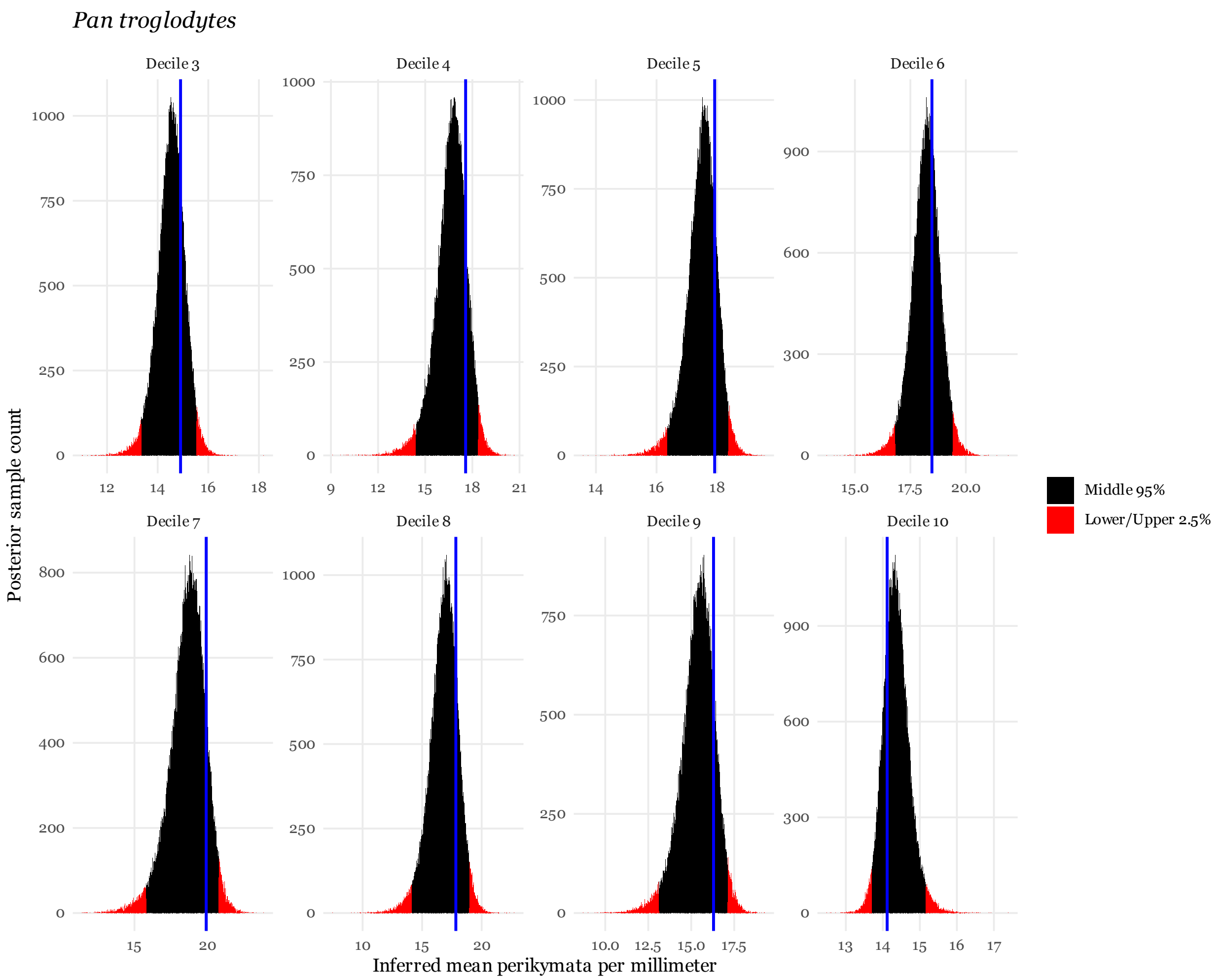


**Figure 25: Posterior distribution for the *Pan troglodytes* intraspecific I² mean.** The horizontal axis indicates the inferred mean perikymata per millimeter for each decile and is truncated between 0 and 30. The vertical axis describes the frequency of that value in the posterior samples. The blue vertical line indicates the empirical mean perikymata per millimeter for each decile. Red bins indicate the lower and upper 2.5% quantiles and the remaining black bins fall within the 95% central credible region. The width of this distribution indicates the statistical uncertainty in the value of this parameter; wider distributions are more uncertain. Histograms were built with 500 bins.

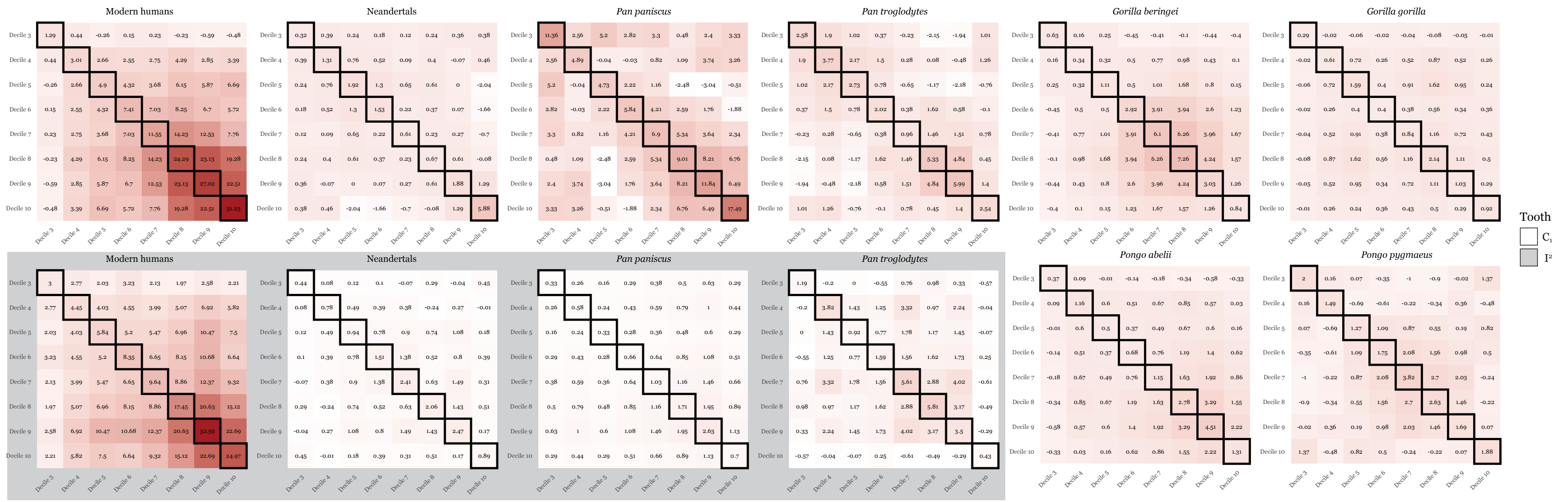


**Figure 26: Posterior mean intraspecific variance-covariance matrices.** Grey show intraspecific VCV matrices from the $I^2$ analysis while all other intraspecific VCV matrices are in white. Diagonal elements indicate the variance in that decile and are bordered with black. Off diagonal elements indicate the covariance across deciles. Darker colors indicate higher rates or covariances. Each mean variance-covariance matrix was calculated on the SPD manifold.

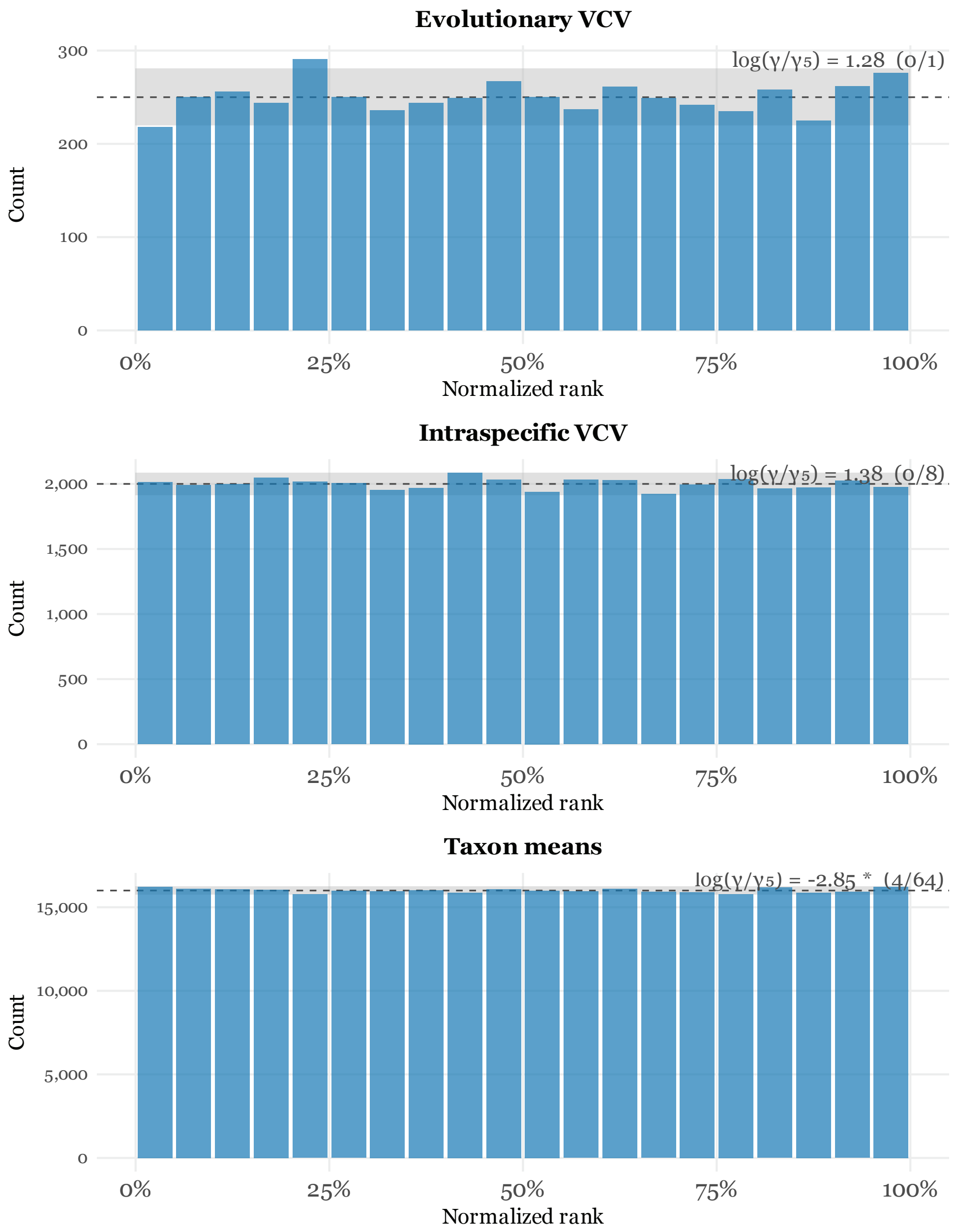


**Figure 27: Simulation study results.** Rank uniform tests for all the evolutionary VCV trace, intraspecific VCV trace (pooled across taxa for visualization), and taxon means (pooled across taxa for visualization) from 5,000 replicates with eight taxa, eight traits, no missing observations, and 2 individuals per taxon. Taxon means are ranked on a per-element basis. $\log(\gamma/\gamma_5)$ in the top right of each histogram indicates the value of the lowest $\log(\gamma/\gamma_5)$ of any parameter in that category and the number of parameters in that category that deviate significantly from uniformity.

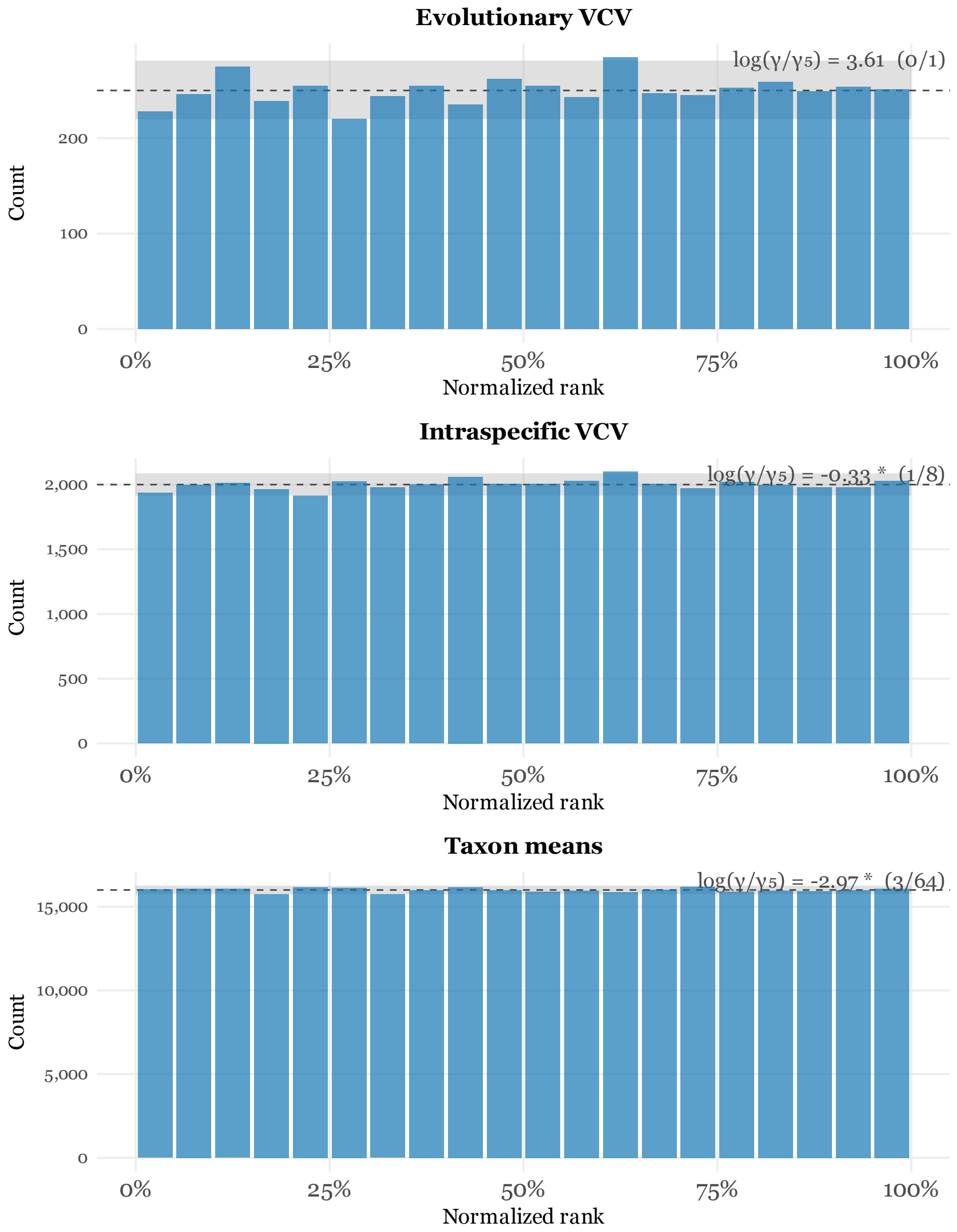


**Figure 28: Simulation study results.** Rank uniform tests for all the evolutionary VCV trace, intraspecific VCV trace (pooled across taxa for visualization), and taxon means (pooled across taxa for visualization) from 5,000 replicates with eight taxa, eight traits, no missing observations, and 16 individuals per taxon. Taxon means are ranked on a per-element basis. $\log(\gamma/\gamma_5)$ in the top right of each histogram indicates the value of the lowest $\log(\gamma/\gamma_5)$ of any parameter in that category and the number of parameters in that category that deviate significantly from uniformity.

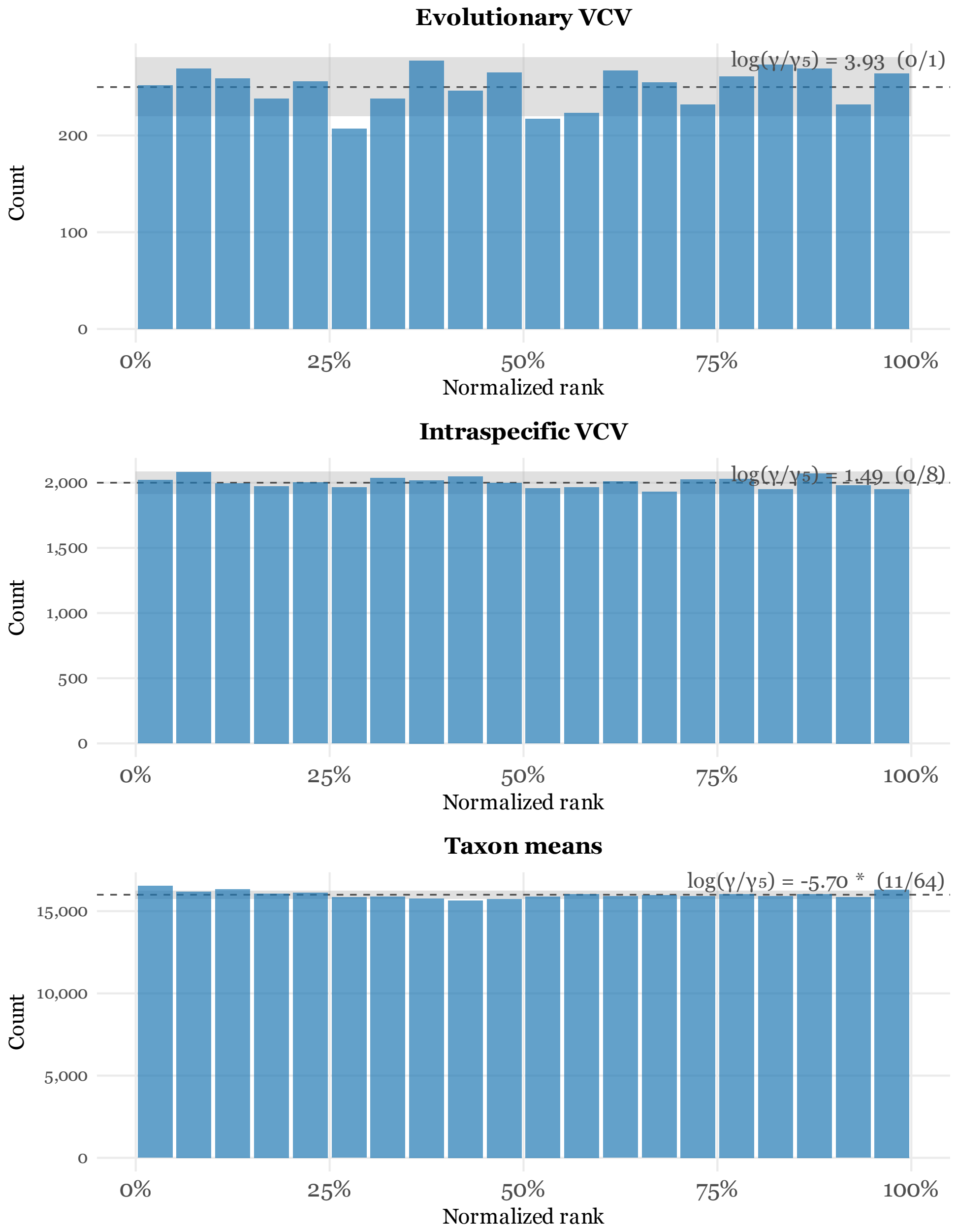


**Figure 29: Simulation study results.** Rank uniform tests for all the evolutionary VCV trace, intraspecific VCV trace (pooled across taxa for visualization), and taxon means (pooled across taxa for visualization) from 5,000 replicates with eight taxa, eight traits, 32 missing observations, and 2 individuals per taxon. Taxon means are ranked on a per-element basis. $\log(\gamma/\gamma_5)$ in the top right of each histogram indicates the value of the lowest $\log(\gamma/\gamma_5)$ of any parameter in that category and the number of parameters in that category that deviate significantly from uniformity.
